# A hypothalamic dynorphin-KOR pathway that orchestrates male sexual behavior under sexual competition

**DOI:** 10.64898/2026.09.01.748525

**Authors:** Quanchi Lyu, Yang Zheng, Xun Ma, Luping Yin

**Affiliations:** Westlake Laboratory of Life Sciences and Biomedicine, Hangzhou 310024, China; School of Life Sciences, Westlake University, Hangzhou 310024, China; Institute of Biology, Westlake Institute for Advanced Study, Hangzhou 310024, China

**Keywords:** sexual competition, aggression-mating trade-off, dynorphin, kappa-opioid receptor, MPOA, VMHvl, Arc, neural plasticity

## Abstract

In nature, males often attack rivals before mating, yet how aggression shapes mating strategies remains unclear. Using a laboratory sexual competition model with optogenetics, calcium imaging, and pharmacology, we found that aggressive male mice attack competitors before mating, whereas social males mate promptly. This delay is mediated by Gi-coupled kappa-opioid receptor-expressing GABAergic interneurons in the medial preoptic area (MPOA^KOR^), recruited by estrogen receptor 1-expressing neurons in the ventrolateral subdivision of the ventromedial hypothalamus (VMHvl^Esr1^), which inhibit MPOA^Esr1^ mating neurons. VMHvl^Esr1^ neurons activate MPOA^KOR^ neurons through direct glutamatergic transmission and indirect disinhibition by suppressing dynorphin release from prodynorphin-expressing arcuate nucleus terminals (Arc^Pdyn^). During rival encounters, aggressive males show a greater MPOA dynorphin decline and stronger MPOA^KOR^ activity than social males. Thus, a VMHvl^Esr1^/Arc^Pdyn^→MPOA^KOR^→MPOA^Esr1^ circuit gates mating according to prior aggressive experience, explaining how social competition shapes reproductive strategies.

**HIGHLIGHTS:**

- Males with distinct aggression levels exhibit divergent reproductive strategies: aggressive males compete before mating; non-aggressive social males mate promptly.
- MPOA^KOR^ interneurons act as the long-sought relay, mediating inhibition from VMHvl^Esr1^ to MPOA^Esr1^ neurons to delay mating.
- VMHvl glutamatergic excitation and Arc dynorphin disinhibition collectively activate MPOA^KOR^ neurons.
- Aggressive males sustain competitor-induced MPOA dynorphin suppression to maintain MPOA^KOR^ in high activity during aggression training.

## INTRODUCTION

To maximize reproductive fitness, animals evolve context-dependent mating strategies shaped by social hierarchy and environmental cues^1,2^. Such adaptive reproductive behaviors are primarily driven by two canonical modes of sexual selection: intrasexual competition and intersexual mate choice. Mammals, including most primates and rodents, predominantly rely on intrasexual competition, whereby males engage in conspecific aggression to secure mating precedence. Birds and fish, however, tend to prioritize intersexual mate choice, with females selecting mates based on male ornamental traits and ritualized courtship performances. The former intrasexual competitive reproductive strategy is widely conserved across animal taxa and is well-characterized in mice^3–5^. In the wild, male mice integrate social competition and mating cues to make adaptive reproductive decisions within complex multi-animal groups. However, most neurobiological studies rely on simplified two-animal tests in laboratory settings. While these minimalistic assays have successfully uncovered core social neural circuits, they fail to recapitulate natural competitive social contexts, leaving the neural mechanisms underlying context-dependent mating regulation during male competition poorly understood.

Existing neurobiological studies have delineated the neural circuits governing aggression and mating primarily via two-animal behavioral assays, establishing a foundational framework for dissecting the neural mechanisms governing reproductive decision-making in natural competitive contexts. In mammals, the hypothalamus orchestrates these two competing innate behaviors through distinct neuronal subpopulations^6^. Glutamatergic estrogen receptor 1-expressing neurons in the ventrolateral subdivision of the ventromedial hypothalamus (VMHvl^Esr1^) neurons drive innate aggression: their activation triggers attacks, whereas their inhibition suppresses such behavior^7^. In contrast, GABAergic Esr1-expressing neurons in the medial preoptic area (MPOA ^Esr1^) neurons are indispensable for the initiation of male mating behavior^8,9^. The VMHvl and MPOA form reciprocal regulatory connections to balance social behavioral outputs: MPOA-to-VMHvl inhibitory signaling restrains aggression toward dominant opponents^10^, whereas VMHvl-to-MPOA signaling inhibits mating attempts toward females in males^8^. Given the intrinsic excitatory property of VMHvl^Esr1^ neurons^11,12^, it remains a long-standing mechanistic paradox how excitatory VMHvl inputs inhibit mating-promoting MPOA^Esr1^ activity. Despite extensive efforts, the neuronal substrate and anatomical location of this critical suppressive relay have never been definitively identified. Of note, the MPOA, though traditionally recognized as a dedicated mating-regulating center, exhibits prominent cellular heterogeneity, as uncovered by recent single-cell sequencing and MERFISH analyses. These high-resolution profiling studies have identified distinct MPOA inhibitory subtypes, including *Oprk1^+^*, *Th*^+^, and *Cck*^+^ neurons, which are specifically activated during male competitive aggression^13^. These aggression-responsive inhibitory populations provide a plausible mechanistic clue to resolve the longstanding VMHvl–MPOA inhibitory paradox.

This persistent circuit ambiguity stems largely from insufficient characterization of hypothalamic local microcircuits. Single-cell transcriptomic and spatial profiling has uncovered extensive neuronal diversity within the MPOA. However, the functions of most molecularly defined neuronal populations and their internal circuit connectivity remain poorly resolved^13^. Recently, large-scale paired whole-cell recordings across hundreds of neuronal pairs have confirmed that individual hypothalamic nuclei exhibit extremely sparse intranuclear synaptic connectivity, whereby local neural communication depends primarily on neuropeptidergic signaling rather than fast chemical synaptic transmission^14–16^. This wiring scheme differs fundamentally from the densely synaptic, rapidly responsive cortical circuitry specialized for fast sensory and motor processing^17^. The neuropeptide-dominated hypothalamic architecture is optimized for slow, sustained physiological transitions, including hunger, temperature homeostasis, maternal care, and mating, yet it cannot account for the rapid, second-scale behavioral switching between aggression and mating during male competitive interactions. Such fast context-dependent social decision-making presumably requires dynamic fast synaptic communication among molecularly distinct MPOA neuronal subsets. To date, however, the local MPOA microcircuits that mediate this rapid competitive behavioral selection remain unidentified.

Here, we established a sexual competition paradigm to explore circuit mechanisms governing male mating-aggression choices amid male and female social partners. We found that aggressive males prioritize rival attack and delay mating initiation, whereas social males remain unaffected. We identified a population of Gi-coupled kappa-opioid receptor (KOR)-expressing neurons in the MPOA (MPOA^KOR^) as an inhibitory relay bridging VMHvl^Esr1^ and MPOA^Esr1^ neurons. During male–male competition, VMHvl^Esr1^ neurons modulate KOR neuron activity via dual pathways: direct glutamatergic excitation and indirect reduction of arcuate nucleus (Arc)-derived dynorphin release. This dynorphin-KOR axis exhibits experience-dependent plasticity: social males habituate to competitor-induced dynorphin decline, whereas aggressive males sustain persistent dynorphin reduction to maintain elevated MPOA^KOR^ excitability. We thus define a hypothalamic circuit: VMHvl^Esr1^/Arc^Pdyn^→MPOA^KOR^→MPOA^Esr1^. This KOR-mediated suppression of reproductive circuitry enables the brain to “compete first and mate later” during sexual competition, elucidating a neural strategy underlying the evolutionary trade-off between competitive dominance and reproductive success.

## RESULTS

### Sexual competition delays mating initiation in aggressive males but not in social males

To uncover the neural mechanisms underlying context-dependent adaptive reproductive decisions in a more natural competitive environments, we established a three-animal sexual competition paradigm that better approximates ethological social contexts compared with conventional two-animal assays. Here, we sought to examine how a male’s aggression level influences its mating strategy when confronted with both a male competitor and a female. Before testing, sexually experienced C57BL/6J male mice were single-housed for three days to establish territory and enhance aggression. A receptive female was introduced for a regular sexual behavior test (RS test) to assess baseline mating behavior. Subsequently, a BALB/c male was introduced for 10 min/day over five days. Based on their behavioral responses to the intruder, males were dichotomized into social (only contact) and aggression groups. Following training, males underwent a sexual competition test (SC test), in which a receptive female and a BALB/c male were simultaneously introduced (Fig. 1A). Notably, no mating between BALB/c males and females was observed (0 of the total 38 tests in this research); all mating was performed by C57BL/6J males.

**Figure 1.**
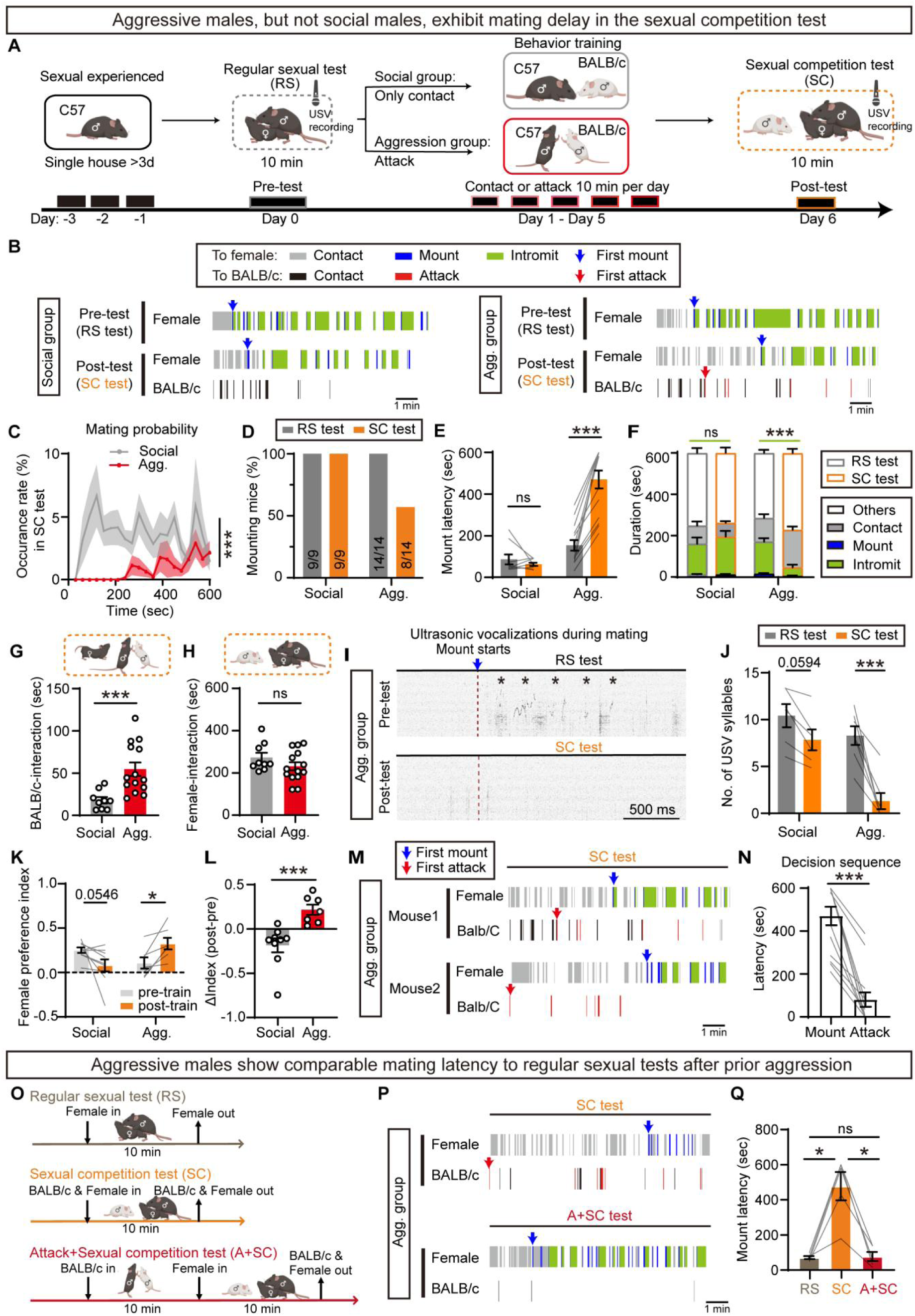
Aggressive males exhibit delayed sexual behaviors during sexual competition. (A) Experimental timeline of behavioral test. (B) Behavioral raster plots of a social male and an aggressive male in pre-test (regular sexual test, RS test) and post-test (sexual competition test, SC test) toward the receptive female or the intruder BALB/c male. (C) Mating probability of social group or aggression group in sexual competition test. (D-F) Percentage of mating mice during the 10-min test (D), mount latency (E), and behavior distribution (contact, mount, and intromit) (F) within social and aggression group. (G-H) Total interaction time with BALB/c males (contact and attack) (G) and total interaction time with females (contact, mount, and intromit) (H) during the sexual competition test. (I) Example ultrasonic vocalization (USV) spectrogram of aggression group during mounting in RS test and SC test. Asterisks indicate USV syllables. (J) Number of USV during each mounting episode. (K-L) Female-preference index of social and aggression group pre- and post-training (K), and the change of index (L). (M) Behavioral raster plots of aggression group in sexual competition test toward the female or BALB/c. Blue and red arrows represent the first mount and first attack, respectively. (N) Latency of mount and attack of aggression group in sexual competition test. (O-Q) Experimental design for the paradigm with 10-min prior BALB/c male exposure before female introduction (O, A+SC test); representative behavioral raster plots (P); and mount latency (Q) across different experimental conditions in aggression group males. Data are mean±SEM. (C) Two-way ANOVA. (E, J, K) Two-way repeated measures ANOVA with Sidak’s multiple comparisons test. (F) Two-way repeated measures ANOVA of intromission duration, followed by Sidak’s multiple comparisons test. (G) Welch’s unpaired t-test. (H) Two-tailed unpaired t test. (L) Two-tailed Mann-Whitney test. (N) Two-tailed Wilcoxon matched-pairs signed rank test. (Q) One-way repeated-meassures ANOVA followed by Tucky’s multiple comparisons test. \**p* < 0.05, \*\**p* < 0.01, \*\*\**p* < 0.001; ns, not significant. n=animals. (C-H) n=9 in social group and 14 in aggression group. (J) n=6 in social group and 7 in aggression group. (K, L) n=9 in social group and 7 in aggression group. (N) n=14. (Q) n=5. See also Figure S1.

In the social group, the BALB/c competitor did not significantly alter mating performance versus RS pre-test; all males mated within the 10-min SC test with no change in mount latency or time spent mating, indicating the competitor does not interfere with mating in social males (Fig. 1B-1F and S1A-S1C). In contrast, the aggression group showed a markedly different profile: the competitor significantly delayed sexual initiation, accompanied by prominent attacks toward BALB/c (Fig. 1B-1C). Only 57% (8/14) of aggression group mated within 10 min (Fig. 1D), with prolonged mount latency and reduced mating time (Fig. 1E-1F), indicating that aggression disrupted mating progression. Time allocation analysis revealed that aggression group spent more time interacting with BALB/c than social group, whereas time with the female did not differ between groups (Fig. 1G-1H), suggesting that mating delay was not due to reduced female contact but rather that aggression blocked conversion of contact into mounting. Among aggression groups that mated, intromission duration and inter-intromission intervals were comparable to pre-test and social group values (Fig. S1A-S1B), indicating the competitor impaired mating initiation, not sexual capacity. Temporal analysis showed that social males initiated mating within ∼200 s and maintained stable copulation, whereas contact with BALB/c declined over time, whereas aggression group sustained high levels of contact and attack throughout, with only a subset transitioning to mating mid-to-late (Fig. S1C-S1D). During mounting, males emit ultrasonic vocalizations (USVs) to attract females; the competitor reduced USV syllables in both groups, but the reduction was significantly greater in the aggression group (Fig. 1I-1J), indicating that the competitor more substantially attenuates mating drive in aggressive males.

The behavioral differences between the social and aggression groups during the SC test might reflect differences in social preference toward females versus BALB/c males. Before training, both groups showed comparable preference to the female. After training, the female preference index of the social group declined, whereas that of the aggression group increased (Fig. 1K-1L), suggesting that aggressive males develop heightened female preference following competitor exposure, reinforcing perception of BALB/c as a rival and contributing to mating delay. Aggression group typically attacked BALB/c before mating during SC test (Fig. 1M-1N), suggesting that mating delay reflects a strategy of first establishing dominance before mating. To test this, aggression groups were first co-housed with BALB/c for 10 min to establish dominance, followed by introduction of a receptive female (Fig. 1O); prior attack significantly shortened mount latency to pre-test levels (Fig. 1P-1Q), supporting that mating delay arises from the need to first establish dominance. To test whether the mating delay requires sexual competition specifically rather than attack itself, we substituted juvenile males (P15–20) for BALB/c (Fig. S1E); although males still attacked juveniles, no mating delay was observed—mount latency, temporal distribution, and USV syllables were comparable to pre-test (Fig. S1F-S1I). Together, these findings demonstrate that a sexual competitor specifically induces mating delay in aggressive males, while leaving social males unaffected. Aggressive males preferentially attack the competitor before mating, reflecting both a stronger drive toward the female and a strategic requirement to first establish competitive dominance for reproductive success.

### VMHvl^Esr1^ neurons send indirect inhibitory projections to MPOA^Esr1^ neurons

We next investigated the neural mechanisms through which aggressive experience modulates male mating strategies. We hypothesized that aggression-associated VMHvl neurons inhibit MPOA circuits that promote mating, thereby mediating delayed mating behavior in aggressive males. To test this, we injected a retrograde tracer (AAV11-fDIO-Cre-GFP) into the MPOA of Vglut2-Flp or Vgat-Flp male mice to label its glutamatergic or GABAergic inputs (Fig. 2A). In Vglut2-Flp mice, abundant GFP^+^ neurons were identified in the VMHvl (Fig. 2B-2C), establishing the VMHvl as a major glutamatergic input source to the MPOA. By contrast, no GFP⁺ neurons were detected in or around the VMHvl in Vgat-Flp mice (Fig. 2B-2C). Thus, under our tracing conditions, the VMHvl-to-MPOA projection was overwhelmingly glutamatergic, with no detectable contribution from GABAergic neurons in the VMHvl. Immunohistochemistry revealed that ∼85% of glutamatergic VMHvl→MPOA neurons expressed Esr1, and ∼60% of VMHvl^Esr1^ neurons projected to MPOA (Fig. 2B and 2D). Subsequent functional experiments focused specifically on the VMHvl^Esr1^→MPOA projection.

**Figure 2.**
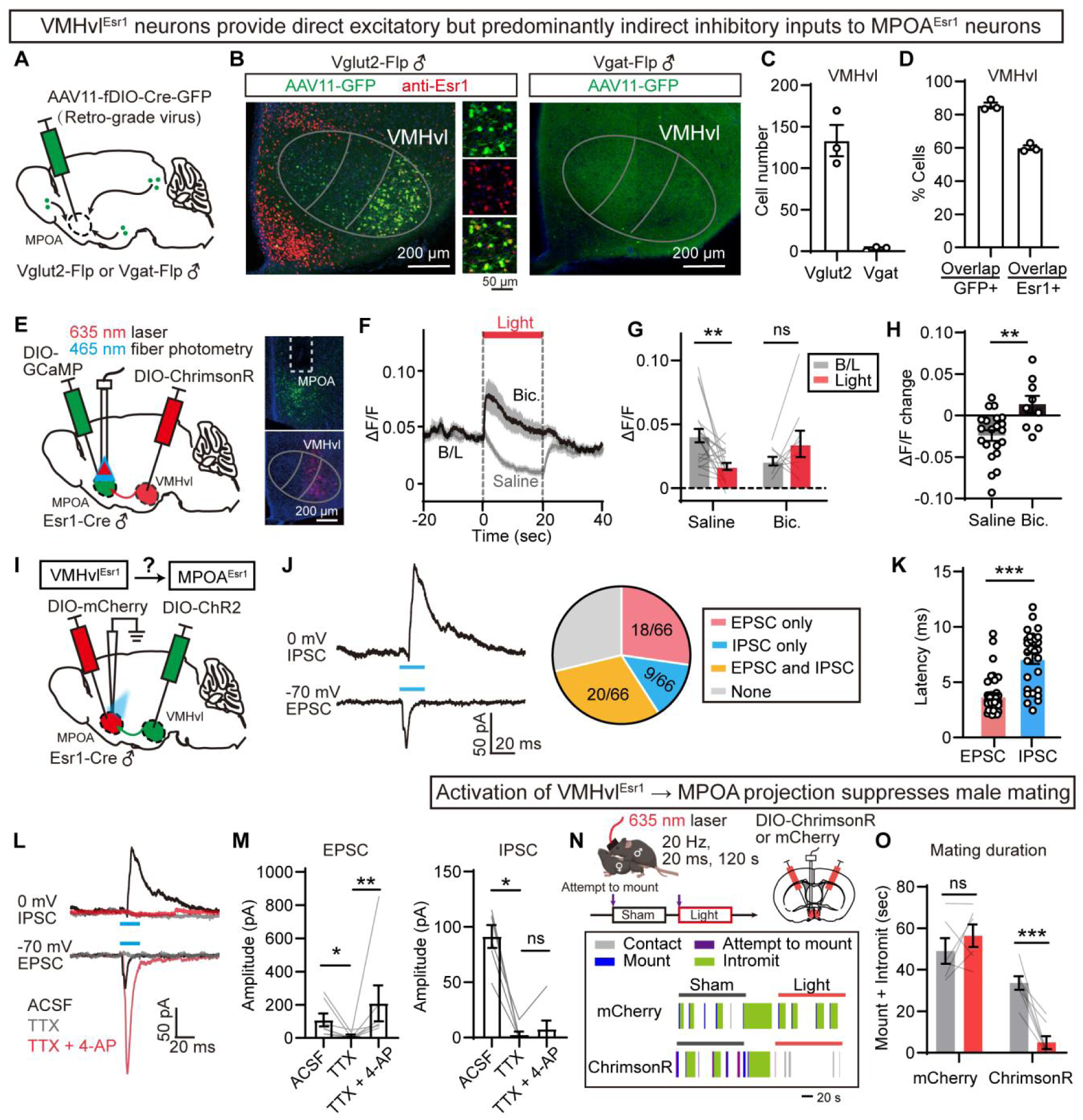
VMHvl^Esr1^ neurons send indirect inhibitory projections to MPOA^Esr1^ neurons. (A) Strategy for labeling glutamatergic (Vglut2-Flp) or GABAergic (Vgat-Flp) upstream neurons projecting to MPOA via AAV11-GFP. (B) Representative VMHvl sections showing AAV11-GFP (green) and Esr1 (red) in Vglut2- Flp (left) and Vgat-Flp (right) mice. (C) Quantification of AAV11-GFP^+^ cells in VMHvl of Vglut2-Flp vs. Vgat-Flp mice. (D) Overlap of AAV11-GFP^+^ and Esr1^+^ cells in VMHvl of Vglut2-Flp mice. (E) Viral strategy and histological validation for simultaneous optogenetic activation of the VMHvl^Esr1^→MPOA projection and GCaMP calcium recording in MPOA^Esr1^ neurons. (F) Averaged GCaMP signals in MPOA^Esr1^ neurons across animals upon light activation of the VMHvl^Esr1^→MPOA projection after i.p. injection of saline or GABA_A_ receptor blocker bicuculline (Bic). B/L, baseline. (G-H) Mean ΔF/F signals at baseline and during light stimulation (G), and the corresponding signal changes (light − B/L) (H) in the saline and bicuculline groups. (I) Viral strategy for recording synaptic connection between VMHvl^Esr1^ and MPOA^Esr1^ in brain slices. (J) Representative IPSC/EPSC traces from MPOA^Esr1^ neurons (left) and pie chart showing the proportion of MPOA^Esr1^ neurons receiving IPSCs or EPSCs (right). (K) Latency of opto-evoked EPSCs and IPSCs. (L-M) Representative IPSC and EPSC traces (L) and amplitude statistics (M) of MPOA^Esr1^ cells after TTX and 4-AP application. (N) Viral strategy for activating the VMHvl^Esr1^→MPOA projection, light stimulation paradigms, and representative behavioral annotations for mCherry and ChrimsonR groups. (O) Mating duration (mount + intromit) during sham and light. Data are mean±SEM. (G, O) Two-way repeated measures ANOVA with Sidak’s multiple comparisons test. (H) Two-tailed unpaired t test. (K) Two-tailed Mann-Whitney test. (M) Friedman test with repeated measures followed by Dunn’s multiple comparisons test. \**p* < 0.05, \*\**p* < 0.01, \*\*\**p* < 0.001; ns, not significant. n=animals; N=cells. (C, D) n=3. (G, H) n=21 for saline group; n=9 for bicuculline group. (K) N=38 MPOA^Esr1^ cells for EPSC; 29 for IPSC. (M, left) N=7 MPOA^Esr1^ cells for EPSC. (M, right) N=6 MPOA^Esr1^ cells for IPSC. (O) n=6 mice for mCherry; n= 7 for ChrimsonR. See also Figure S2.

To test how VMHvl^Esr1^ activation modulates MPOA^Esr1^ activity, we combined optogenetic stimulation of VMHvl^Esr1^ terminals in MPOA with GCaMP imaging of MPOA^Esr1^ neurons (Fig. 2E). VMHvl^Esr1^ terminal activation significantly decreased MPOA^Esr1^ calcium activity, an effect blocked by the GABA_A_ receptor antagonist bicuculline (Fig. 2F-2H). Given that VMHvl^Esr1^ neurons are glutamatergic, this inhibition likely occurs through recruitment of local MPOA interneurons that indirectly suppress MPOA^Esr1^ neurons.

To directly characterize synaptic connectivity, we performed ex vivo patch-clamp recordings from MPOA^Esr1^ neurons while stimulating VMHvl^Esr1^ terminals (Fig. 2I-2J). Of 66 MPOA^Esr1^ neurons, ∼57% (38/66) exhibited EPSCs, 44% (29/66) IPSCs, and 30% (20/66) both. IPSC latency (7.14 ± 0.47 ms) was significantly longer than EPSC latency (3.75 ± 0.29 ms) (Fig. 2J-2K). TTX application reduced EPSC amplitude, which was rescued by 4-AP, indicating monosynaptic direct input; in contrast, IPSCs could not be rescued by 4-AP (Fig. 2L-2M), demonstrating polysynaptic indirect inhibition. Thus, VMHvl^Esr1^ neurons exert monosynaptic excitation and polysynaptic inhibition onto MPOA^Esr1^ neurons, with net inhibitory effect upon activation. MPOA^Tacr1^ neurons, a subpopulation implicated in sexual behavior, showed similar response profiles (EPSCs: 27/34=79%; IPSCs: 24/34=71%) (Fig. S2A-S2D), further supporting that VMHvl^Esr1^ sends convergent direct excitatory and indirect inhibitory projections to MPOA^Esr1^ neurons controlling sexual behavior.

Consistent with the inhibition of MPOA^Esr1^ mating neurons upon VMHvl^Esr1^ activation, optogenetic activation of VMHvl^Esr1^ terminals unilaterally in the MPOA significantly suppressed male sexual behavior (Fig. 2N-2O). To confirm that our unilateral light delivery adequately activated bilateral terminals, we verified that unilateral light delivery to the MPOA was sufficient for bilateral activation of ChrimsonR-expressing neurons, as confirmed by Fos/ChrimsonR overlap, while ruling out trans-hemispheric synaptic transmission (Fig. S2E-S2F). Integrating electrophysiological and behavioral evidence, we propose that during encounters with a sexual competitor, VMHvl^Esr1^ neurons indirectly suppress MPOA^Esr1^ activity via a polysynaptic pathway, thereby delaying mating initiation. This provides a circuit basis for the neural regulation of aggression–sexual behavior interaction.

### Attack-activated MPOA^KOR^ neurons suppress sexual behavior

We hypothesized that a subpopulation of MPOA neurons activated during aggressive behavior—hereafter referred to as “attack neurons”—receives excitatory inputs from VMHvl and suppresses MPOA^Esr1^ neuronal activity via GABA release, thereby delaying sexual behavior. To test this, we employed the TRAP (targeted recombination in active populations) strategy to selectively label attack-activated neurons (TRAP-attack neurons) in the MPOA. Fos-CreER mice were bilaterally injected with AAV-DIO-ChR2-mCherry into the MPOA. Three weeks later, males underwent 5 days of aggression training, with social as control. On the final day, they were subjected to a 15-minute resident-intruder aggression test with a BALB/c intruder, followed immediately by 4-OHT administration to induce permanent ChR2 expression in neurons activated during aggression. Two weeks after labeling, we optogenetically activated these TRAP-labeled neurons and assessed their effects on sexual behavior (Fig. 3A). To validate TRAP labeling efficiency, we exposed mice to a BALB/c intruder 1 hour before perfusion; co-immunostaining for ChR2-mCherry and Fos revealed approximately 50% co-labeled cells (Fig. 3B-3C).

**Figure 3.**
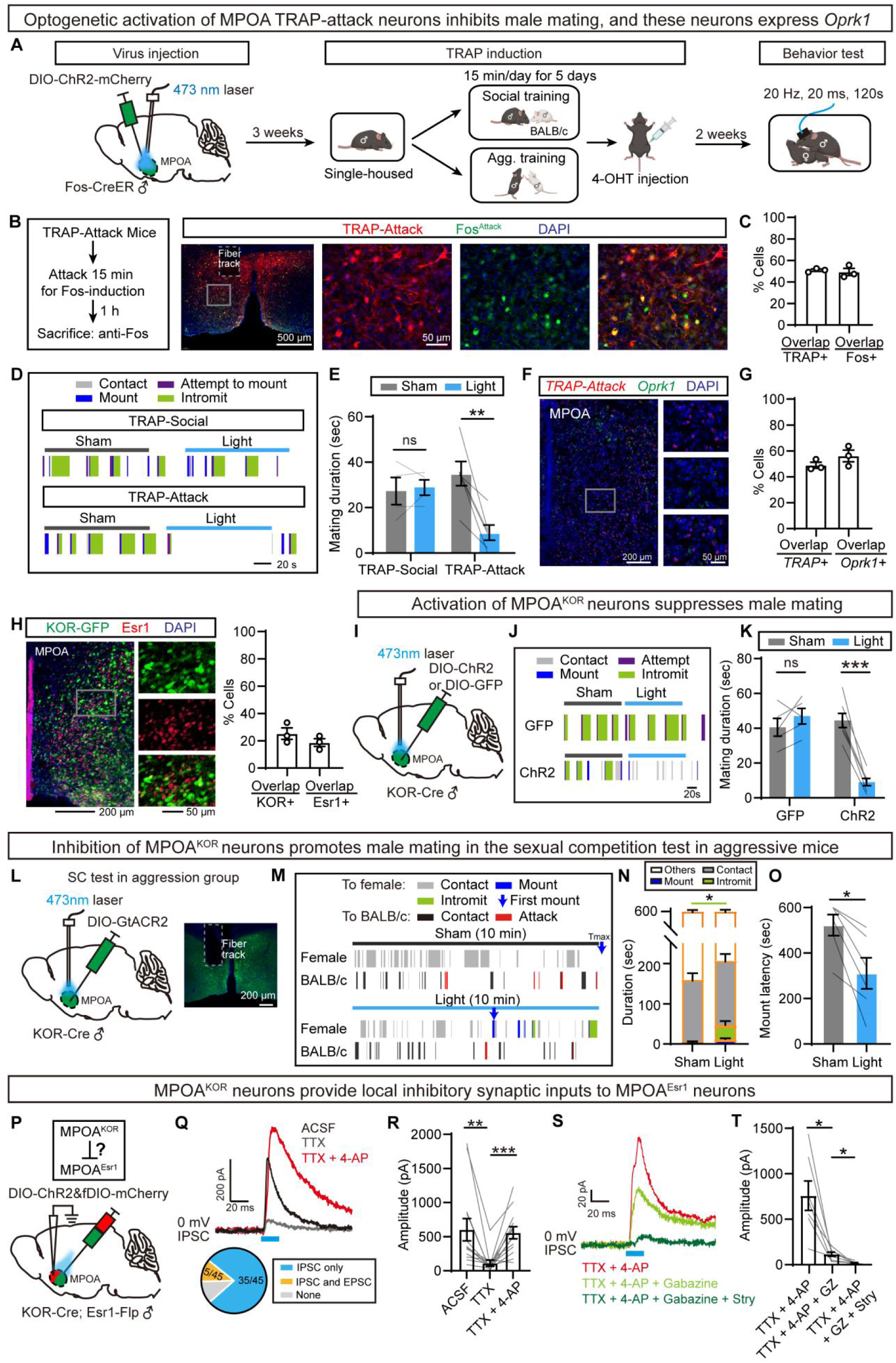
Attack-activated MPOA neurons express KOR and their stimulation suppresses sexual behaviors. (A) Experimental strategy of TRAP for labeling and activating attack-activated neurons in the MPOA. (B) Representative images of TRAP-attack neurons and attack-evoked Fos immunostaining in the MPOA. (C) Overlap between TRAP-attack neurons and attack-induced Fos staining in the MPOA. (D) Behavioral raster plots of Fos-CreER males injecting DIO-ChR2 (social or attack-induced) in the MPOA during female intruder exposure under sham and light simulation. (E) Mating duration upon sham and light simulation of TRAP-social or TRAP-attack neurons. (F) RNAscope of TRAP-attack neurons and *Oprk1* mRNA. (G) Overlap between *Oprk1* mRNA and TRAP-attack neurons . (H) Viral-GFP labeled KOR-positive neurons and Esr1 immunostaining (left), and their colocalization (right). (I) Viral strategy for opto-activating MPOA^KOR^ neurons. (J) Behavioral raster plots of KOR-Cre males injected with DIO-GFP or DIO-ChR2 in the MPOA with a female intruder under sham and light simulation. (K) Mating duration following sham and light activation of MPOA^KOR^ neurons. (L) Viral strategy for opto-inhibiting MPOA^KOR^ neurons. (M) Behavioral raster plots of KOR-Cre males injected with DIO-GtACR2 in the MPOA during sexual competition test under sham and light simulation. Tmax indicates the end of the test. (N-O) Behavior distribution (contact, mount, and intromit) (N) and mount latency (O) following sham and light inhibition of MPOA^KOR^ neurons. (P) Viral strategy for recording synaptic connections between MPOA^KOR^ and MPOA^Esr1^ in brain slices. (Q) Representative IPSC traces recorded with TTX and 4-AP (top) and the proportion of MPOA^Esr1^ neurons receiving IPSCs or EPSCs (bottom). (R) Quantification of IPSC amplitude following TTX and 4-AP application. (S-T) Representative IPSC traces (S) and amplitude quantification (T) under sequential pharmacological treatment: TTX + 4-AP, TTX + 4-AP + Gabazine (GZ), and TTX + 4-AP + Gabazine + strychnine (Stry). Data are mean±SEM. (E, K) Two-way ANOVA with Sidak’s multiple comparisons test. (N, O) Two-tailed Wilcoxon matched-pairs signed rank test. (R) Friedman test with repeated measures followed by Dunn’s multiple comparisons test. (T) One-way rmANOVA followed by Tucky’s multiple comparisons test. \**p* < 0.05, \*\**p* < 0.01, \*\*\**p* < 0.001; ns, not significant. n=animals; N=cells. (C, G, H) n=3. (E) n=4 for TRAP-social; n=6 for TRAP-attack. (N, O) n=6. (K) n=5 for GFP group; n=7 for ChR2 group. (R) N=12 MPOA^Esr1^ cells for adding TTX and 4- AP. (T) N=7 MPOA^Esr1^ cells for further adding Gabazine and Stry. See also Figure S3.

Optogenetic activation of TRAP-labeled attack neurons during mounting attempts significantly suppressed sexual behavior; this effect was not observed in social control groups (Fig. 3D-3E). Additionally, activating these MPOA attack neurons did not induce aggressive behavior or alter contact with BALB/c (Fig. S3A-S3C), suggesting they serve as downstream effectors of VMHvl^Esr1^ neurons in relaying aggression-related signals. These results suggest that TRAP-attack neurons are selectively labeled in aggressive males, and that their activation inhibits MPOA^Esr1^ neurons to suppress mating.

Previous single-cell RNA-sequencing and MERFISH profiling of the hypothalamic preoptic region identified several inhibitory neuronal populations preferentially recruited during intermale aggression, with enriched expression of *Oprk1*, *Cck*, or *Th*^13^. We therefore systematically examined whether neurons recruited during attack expressed these markers. We visualized KOR^+^, CCK^+^, and TH^+^ cells in combination with anti-Fos immunostaining for attack-induced neurons and found that only KOR showed substantial co-localization with Fos^+^ cells (∼40%) in the MPOA, whereas CCK and TH labeled only a small fraction of attack-activated neurons (2% and 5%, respectively) (Fig. S3D-S3E). Based on this selective enrichment, we focused subsequent experiments on MPOA^KOR^ neurons. Given the limited reliability of KOR antibodies for further characterization, we turned to RNAscope with an *Oprk1* probe to profile MPOA^KOR^ neurons. RNAscope confirmed that ∼50% of TRAP-labeled attack neurons were *Oprk1^+^* (Fig. 3F-3G). These *Oprk1^+^* neurons were predominantly inhibitory: approximately 85% expressed the GABAergic marker *Slc32a1* (*Vgat*), whereas a smaller fraction expressed the glutamatergic marker *Slc17a6* (*Vglut2*), including a subset that co-expressed both markers (Fig. S3F-S3G). In addition, only ∼20% of MPOA^KOR^ neurons co-expressed Esr1 (Fig. 3H), indicating that KOR^+^ and Esr1^+^ cells are largely two separate populations.

We then employed KOR-Cre mice to test the role of MPOA^KOR^ neurons in male mating. In contrast to the pro-sexual effects of activating the broader MPOA^Vgat^ or MPOA^Esr1^ populations^9^, optogenetic activation of MPOA^KOR^ neurons during mounting attempts significantly suppressed sexual behavior (Fig. 3I-3K). Conversely, silencing MPOA^KOR^ neuronal activity via GtACR2 in aggressive males during the sexual competition test significantly facilitated sexual behavior, as reflected by an increase in intromission duration and a marked decrease in mount latency (Fig. 3L-3O).

To validate the synaptic connection between MPOA^KOR^ and MPOA^Esr1^ neurons, we performed slice recordings in KOR-Cre;Esr1-Flp mice. AAV-DIO-ChR2 was injected into MPOA to activate KOR neurons, and AAV-fDIO-mCherry was co-injected to label Esr1 neurons for whole-cell patch-clamp recordings (Fig. 3P). Of 45 recorded Esr1 neurons, 40 exhibited IPSCs evoked by KOR neuron activation, demonstrating robust inhibitory connectivity (Fig. 3Q). TTX application substantially reduced IPSC amplitude, which was rescued by 4-AP, confirming monosynaptic inhibition (Fig. 3R). Notably, TTX did not completely abolish IPSCs, suggesting involvement of non-GABAergic transmission^18^. Sequential application of Gabazine (GABA_A_ receptor antagonist) and strychnine (glycine receptor antagonist) in the presence of TTX and 4-AP revealed that the IPSC comprised both GABAergic and glycinergic components (Fig. 3S-3T).

Together, these findings support a model in which MPOA^KOR^ neurons are recruited during aggression to inhibit MPOA^Esr1^ neurons and suppress mating initiation; relieving this inhibition restores sexual behavior.

### Intra-MPOA dynorphin eliminates mating delay in aggressive males

Dynorphin is the endogenous ligand of KOR, a Gi-coupled GPCR that suppresses neuronal activity upon activation. To further determine whether dynorphin-mediated KOR signaling in the MPOA can rescue mating delay in aggressive males, we bilaterally infused dynorphin or saline into the MPOA 10 min prior to the sexual competition test (Fig. 4A). Saline-infused aggressive males showed mating delay, whereas dynorphin infusion abolished this suppression: mount latency and total mating duration both recovered to pre-test levels (Fig. 4B-4D), indicating that silencing MPOA^KOR^ neurons prevents VMHvl^Esr1^-evoked inhibition onto MPOA^Esr1^ via the KOR relay. Notably, intra-MPOA dynorphin did not affect attack latency or duration (Fig. S4A-S4C), suggesting that although KOR neurons are downstream of VMHvl^Esr1^, they do not mediate aggressive behavior per se.

**Figure 4.**
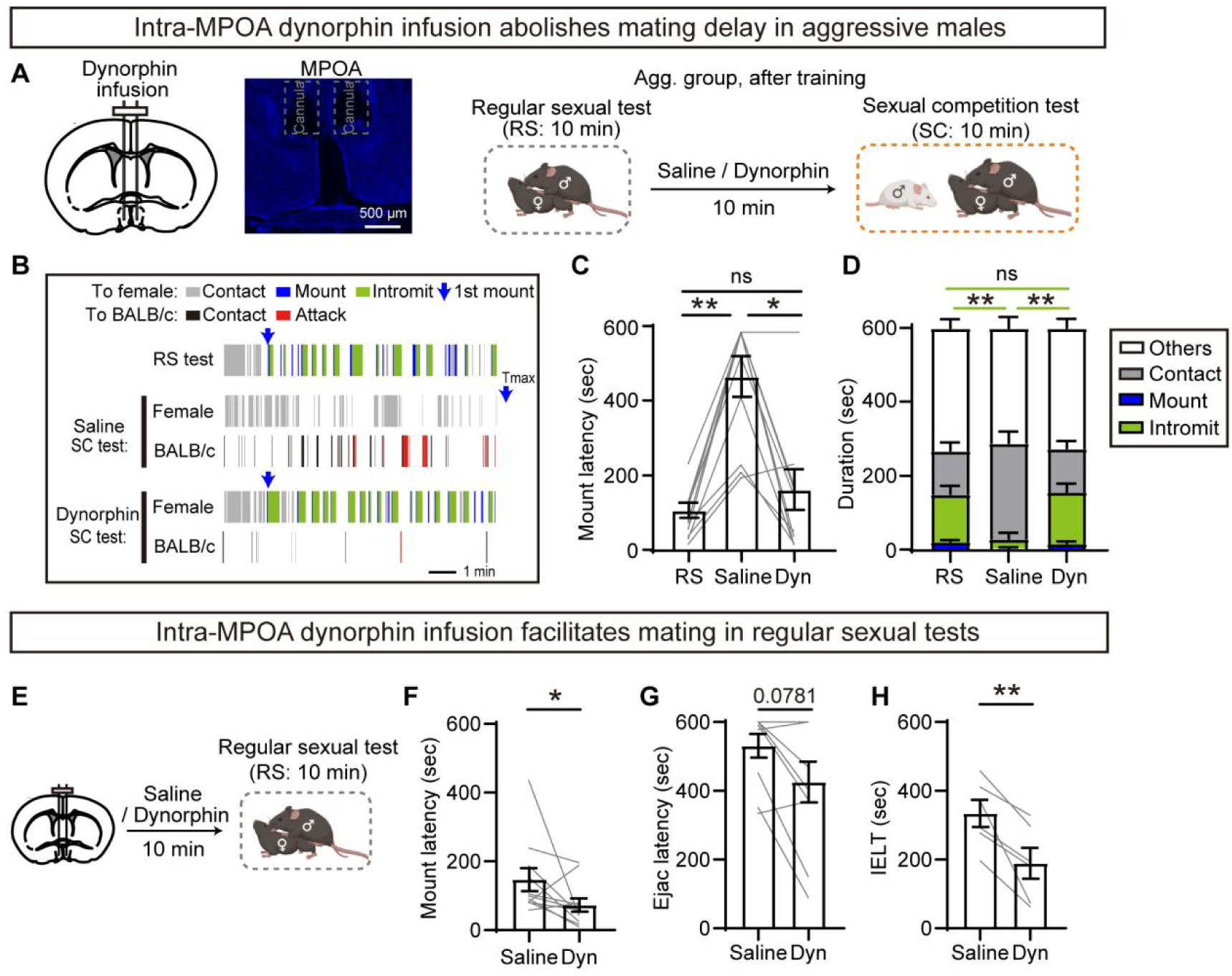
Intra-MPOA dynorphin administration eliminates mating delay in aggressive males. (A) Experimental design of sexual competition test after intra-MPOA dynorphin administration. (B) Behavioral raster plots of an aggression group in RS test, SC test (saline) and SC test (dynorphin). Tmax indicates the end of the test. (C-D) Mount latency (C), and behavior distribution (contact, mount, and intromit) (D) of the aggression group following saline or dynorphin local infusion. (E) Experimental timeline of regular sexual test. (F-H) Changes in mount latency (F), ejaculatory latency (G) and intravaginal ejaculatory latency time (IELT) (H) following intra-MPOA dynorphin administration. Data are mean±SEM. (C, D) Friedman test with repeated measures followed by Dunn’s multiple comparisons test. (F, G) Two-tailed Wilcoxon matched-pairs signed rank test. (H) Two-tailed paired t test. \**p* < 0.05, \*\**p* < 0.01; ns, not significant. n=animals. (C, D, G) n=10. (F) n=11. (H) n=6. See also Figure S4.

Beyond sexual competition, dynorphin infusion also facilitated mating in regular sexual behavior tests, shortening mount latency even without a BALB/c competitor (Fig. 4E-4F), indicating that suppressing MPOA^KOR^ is sufficient to promote sexual initiation independent of VMHvl input. Intra-MPOA dynorphin additionally significantly reduced ejaculatory latency and intravaginal ejaculatory latency time (IELT) (Fig. 4G-4H), which is consistent with previous reports^19^. To pharmacologically activate MPOA^KOR^ neurons in social males, we infused the KOR antagonist nor-BNI into the MPOA (Fig. S4D), predicting that disinhibition would prolong mount latency during the sexual competition test. Contrary to this prediction, nor-BNI treatment did not alter mount latency relative to controls (Fig. S4E). One possibility is that KOR blockade alone is insufficient to drive MPOA^KOR^ neurons to the activity level required to inhibit sexual behavior.

### MPOA^KOR^ neurons relay VMHvl^Esr1^ poly-synaptic inhibition onto MPOA^Esr1^

We have established that MPOA^KOR^ neurons are activated during aggression and regulate sexual behavior via local projections to MPOA^Esr1^ neurons through optogenetics and pharmacological manipulations. To determine that MPOA^KOR^ neurons serve as inhibitory interneurons linking VMHvl^Esr1^ to MPOA^Esr1^, we next performed functional validation both in vivo and in vitro. We first asked whether KOR signaling is required for VMHvl^Esr1^-mediated suppression of MPOA^Esr1^ activity (Fig. 5A). Systemic administration of the KOR agonist U50488 during optogenetic activation of the VMHvl^Esr1^→MPOA projection significantly blocked the decrease in MPOA^Esr1^ calcium signals (Fig. 5B-5D), implicating KOR neurons act as the inhibitory relay. Consistent with this, IPSCs evoked in MPOA^Esr1^ neurons by VMHvl^Esr1^ terminal activation were suppressed by exogenous dynorphin in slice recordings (Fig. 5E-5G). EPSCs in a subset of MPOA^Esr1^ neurons were also modestly reduced (Fig. S5A-S5C), likely reflecting presynaptic KORs on glutamatergic terminals, but the suppression was significantly smaller than that of IPSCs (Fig. S5D). The net effect was disinhibition of MPOA^Esr1^ neurons, supporting MPOA^KOR^ neurons as an indirect inhibitory relay from VMHvl to MPOA^Esr1^. To functionally confirm that MPOA^KOR^ neurons are targets of VMHvl^Esr1^ neurons, we optogenetically stimulated the VMHvl^Esr1^→MPOA projection while monitoring calcium activity in MPOA^KOR^ neurons (Fig. 5H). Terminal stimulation significantly elevated GCaMP signals in MPOA^KOR^ neurons (Fig. 5I-5J), confirming functional recruitment by the VMHvl^Esr1^→MPOA pathway. Corroborating this, optogenetic activation of this projection induced c-Fos expression in the MPOA, with ∼60% of *Fos*^+^ cells co-expressing *Oprk1* (Fig. 5K-5L).

**Figure 5.**
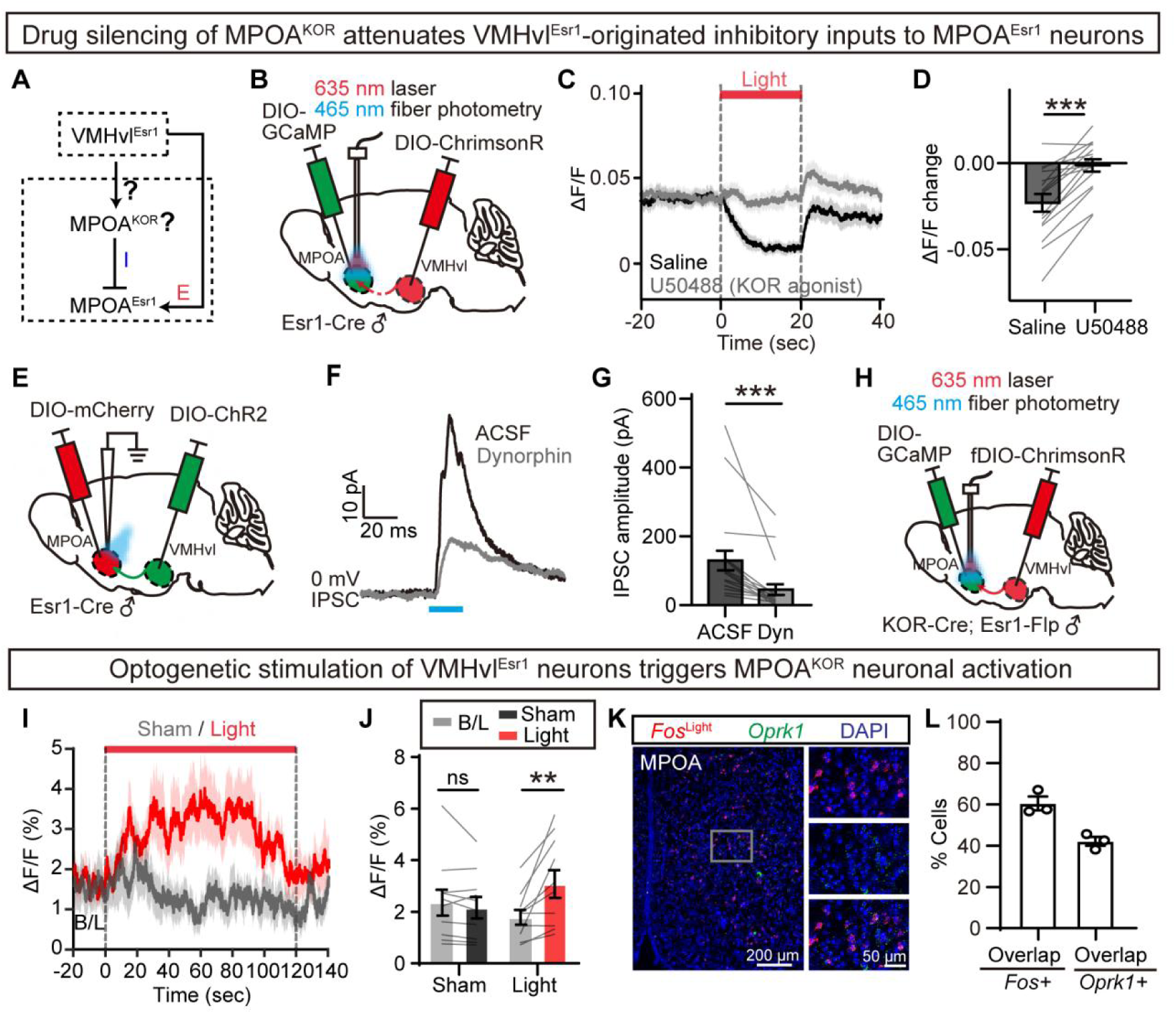
MPOA^KOR^ neurons relay VMHvl^Esr1^ inhibition onto MPOA^Esr1^ neurons. (A) Schematic of the hypothesized circuit of VMHvl^Esr1^→MPOA^KOR^ →MPOA^Esr1^. (B) Viral strategy and histology for opto-activating VMHvl^Esr1^→MPOA projection with simultaneous GCaMP recording in MPOA^Esr1^ neurons. (C) Averaged GCaMP responses in MPOA^Esr1^ across animals during VMHvl^Esr1^→MPOA activation following i.p. injection of saline or U50488. (D) Light-evoked ΔF/F changes (light − baseline) in saline- and U50488-treated groups. (E) Viral strategy for synaptic recording between VMHvl^Esr1^ and MPOA^Esr1^ in brain slices. (F) Representative IPSC traces from MPOA^Esr1^ in ACSF and after dynorphin application. (G) Quantification of IPSC amplitude in MPOA^Esr1^ cells following dynorphin treatment. (H) Viral strategy and histological validation for light-activating VMHvl^Esr1^→MPOA projection with simultaneous GCaMP recording in MPOA^KOR^ neurons. (I) Averaged GCaMP responses in MPOA^KOR^ neurons across animals during VMHvl^Esr1^→MPOA projection activation. B/L, baseline. (J) Mean ΔF/F signal at baseline (B/L) and under sham / light stimulation. (K) RNAscope of light-induced *Fos* and *Oprk1* mRNA in the MPOA. (L) Overlap between light-induced *Fo*s and *Oprk1* in the MPOA. Data are mean±SEM. (D) Two-tailed paired t test. (G) Two-tailed Wilcoxon matched-pairs signed rank test. (J) Two-way ANOVA with Sidak’s multiple comparisons test. \*\**p* < 0.01, \*\*\**p* < 0.001; ns, not significant. n=animals; N=cells. (D) n=16. (G) N=20 MPOA^Esr1^ cells for IPSC. (J) n=10. (L) n=3. See also Figure S5.

### Aggression prevents MPOA^KOR^ habituation to male contact by elevating its intrinsic excitability

Having established that MPOA^KOR^ neurons mediate the inhibitory pathway from VMHvl^Esr1^ to MPOA^Esr1^, we next asked whether the recruitment and plasticity of MPOA^KOR^ neurons differ between aggressive and social males. To investigate MPOA^KOR^ neuronal dynamics during five-day training, we performed GCaMP recordings in social and aggression groups (Fig. 6A). In naive males, both social contact with and attack toward BALB/c reliably elevated KOR calcium activity. Following training, the groups diverged: social group showed significant attenuation of BALB/c-evoked signals, whereas aggression group maintained robust KOR activation during attack (Fig. 6B-6F). This indicates that repeated social exposure without aggression induces habituation of MPOA^KOR^ responsiveness, relieving inhibition onto MPOA^Esr1^ and permitting normal mating; in contrast, sustained aggression maintains persistent KOR activation and delays mating. To isolate neural responses to the same stimulus across groups, we head-fixed males and presented an anesthetized BALB/c male for passive chemosensory investigation (Fig. S6A-S6B). Social group showed significant attenuation in contact-evoked KOR signals across training, whereas aggression group showed no change; post-training KOR activity was significantly higher in aggression than social group (Fig. S6C-S6E), indicating that aggressive experience consolidates MPOA^KOR^ responsiveness to BALB/c cues regardless of contact or attack, sustaining suppression of sexual behavior. Notably, naive social males also showed KOR activation during BALB/c contact before training (Fig. 6E), which was consistent with our finding that introducing a female with a BALB/c male before training prolonged mount latency and reduced mating time even without attack (Fig. S6F-S6I), suggesting that BALB/c presence can delay mating initiation prior to social adaptation.

**Figure 6.**
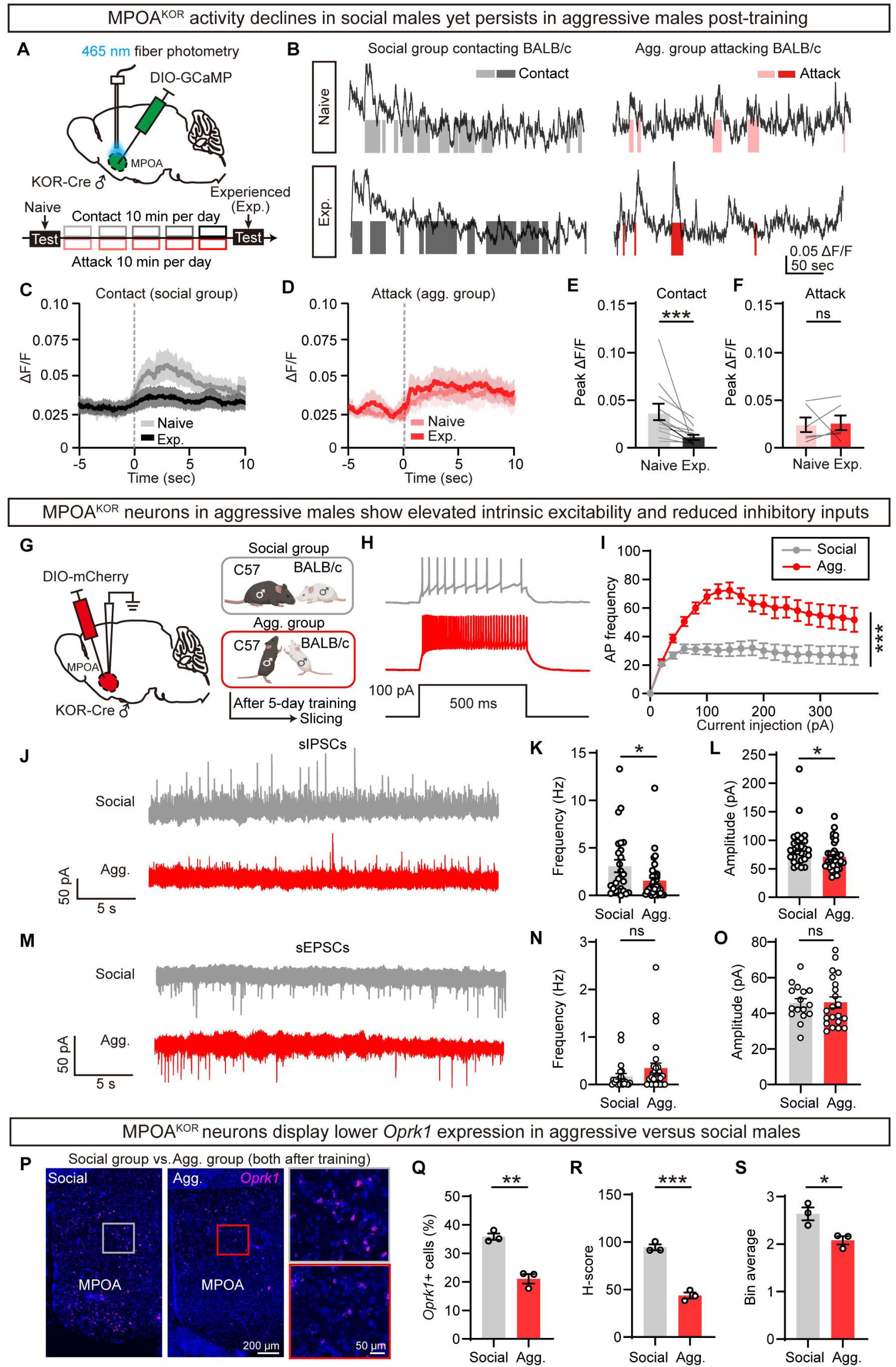
Aggression prevents MPOA^KOR^ adaptation to male contact by elevating its intrinsic excitability. (A) Viral strategy for Ca^2+^ imaging of MPOA^KOR^ neurons (top), and experimental timeline for behavioral tests in social and aggression group (bottom). (B) Example of Ca^2+^ traces from naive and experienced social/aggressive mice. (C) Averaged PETHs of ΔF/F across animals aligned to the onset of contact toward BALB/c in naive (gray) and experienced (black) social mice. (D) Averaged PETHs of ΔF/F across animals aligned to the onset of attack toward BALB/c in naive (pink) and experienced (red) aggressive mice. (E) Peak ΔF/F responses during contacting BALB/c in naive and experienced social mice. (F) Peak ΔF/F responses during during attacking BALB/c in naive and experienced aggressive mice. (G) Experimental strategy for slice recording of MPOA^KOR^ neurons in social and aggression group. (H) Representative action potential traces by 100 pA current injection. (I) F-I curves of MPOA^KOR^ neurons in social and aggression group. (J) Representative traces of spontaneous IPSCs. (K-L) Quantification of sIPSC frequency (K) and amplitude (L). (M) Representative traces of spontaneous EPSCs. (N, O) Quantification of sEPSC frequency (N) and amplitude (O). (P) RNAscope of *Oprk1* mRNA in the MPOA of social and aggressive mice after training. (Q-S) *Oprk1*-positive neuronal proportion (Q), *Oprk1* expression H-score (R), and binned average signal intensity (S) in the MPOA. Data are mean±SEM. (E) Two-tailed Wilcoxon matched-pairs signed rank test. (F) Two-tailed paired t test. (I) Two-way ANOVA. (K, L, N, O) Two-tailed Mann-Whitney test. (Q, R, S) Two-tailed unpaired t test. \**p* < 0.05, \*\**p* < 0.01, \*\*\**p* < 0.001; ns, not significant. n=animals; N=cells. (E) n=12. (F) n=6. (I) N=38 MPOA^KOR^ cells for social group; 37 for aggression group. (K, L) N=26 MPOA^KOR^ cells for social group; 38 for aggression group. (N) N=22 MPOA^KOR^ cells for social group; 28 for aggression group. (O) N=16 MPOA^KOR^ cells for social group; 21 for aggression group. (Q, R, S) n=3. See also Figure S6.

To investigate mechanisms underlying these divergent responses, we next characterized electrophysiological properties of MPOA^KOR^ neurons using whole-cell patch-clamp recordings (Fig. 6G). F-I curves revealed that KOR neurons from aggression group fired at significantly higher rates than those from social group (Fig. 6H-6I), indicating enhanced intrinsic excitability. Analysis of sEPSCs and sIPSCs showed that KOR neurons in the aggression group received sIPSCs of significantly lower frequency and amplitude compared to the social group, while sEPSCs were comparable between groups (Fig. 6J-6O), indicating reduced spontaneous inhibitory input onto KOR neurons in aggression group.

Given that KOR is an inhibitory Gi-coupled receptor, the lower activity state of MPOA^KOR^ neurons in social males may result from elevated Oprk1 expression. To test this, we compared Oprk1 mRNA levels in the MPOA between groups using RNAscope (Fig. 6P). Brains were collected the next day after five-day training, and sections were analyzed for *Oprk1*-positive cell number, H-score, and per-cell expression intensity (Fig. 6Q-6S). Across all measures, the social group showed significantly higher *Oprk1* expression than the aggression group, contributing to the lower neural activity.

Integrating these findings, MPOA^KOR^ neurons in aggression group exhibit greater overall activity towards BALB/c, lower *Oprk1* expression, higher intrinsic excitability, and reduced inhibitory input. This heightened activity strengthens inhibition of MPOA^Esr1^, delaying mating initiation. Together, these results demonstrate that multi-day training produces divergent MPOA^KOR^ plasticity: repeated aggression consolidates MPOA^KOR^ activation to BALB/c cues, whereas social contact habituation attenuates it, providing a circuit mechanism for experience-dependent regulation of mating.

### VMHvl^Esr1^ activates MPOA^KOR^ primarily through dynorphin-reduction-mediated disinhibition

To directly examine how dynorphin–KOR signaling regulates MPOA^KOR^ neuronal activation during sexual competition, we expressed the recently developed dynorphin sensor AAV-kLight1.2 in the MPOA of Esr1-Flp males, together with AAV-fDIO-ChrimsonR delivered to VMHvl for optogenetic activation of the VMHvl^Esr1^→MPOA projection (Fig. 7A). Pathway activation produced a significant decrease in kLight signal (Fig. 7B-7C), indicating reduced extracellular dynorphin, which would disinhibit MPOA^KOR^ neurons. To confirm that the VMHvl-evoked increase in KOR neuron calcium activity is primarily driven by this dynorphin decrease, we performed acute slice electrophysiology (Fig. 7D). Optogenetic activation of the VMHvl^Esr1^→MPOA projection significantly increased MPOA^KOR^ neuron firing, an effect that was abolished by the KOR antagonist nor-BNI (Fig. 7E-7G). These findings point to a disinhibitory mechanism, wherein VMHvl^Esr1^ activity elevates KOR neuron excitability primarily by reducing local dynorphin levels rather than via direct glutamatergic transmission. To further determine whether VMHvl^Esr1^ neurons form direct monosynaptic connections with MPOA^KOR^ neurons, we injected a retrograde transsynaptic rabies virus into the MPOA of KOR-Cre mice (Fig. S7A). This revealed that only ∼20% of the MPOA^KOR^-projecting neurons in the VMHvl expressed Esr1 (Fig. S7B-S7C), indicating that Esr1^+^ inputs represent a minority of total VMHvl afferents to the MPOA^KOR^. Despite this anatomical sparsity, the VMHvl^Esr1^-to-MPOA^KOR^ direct glutamatergic projection is required to drive the poly-synaptic IPSCs from VMHvl^Esr1^ to MPOA^Esr1^, suggesting that this minority subpopulation exerts a disproportionately potent functional influence over MPOA^KOR^ activity.

**Figure 7.**
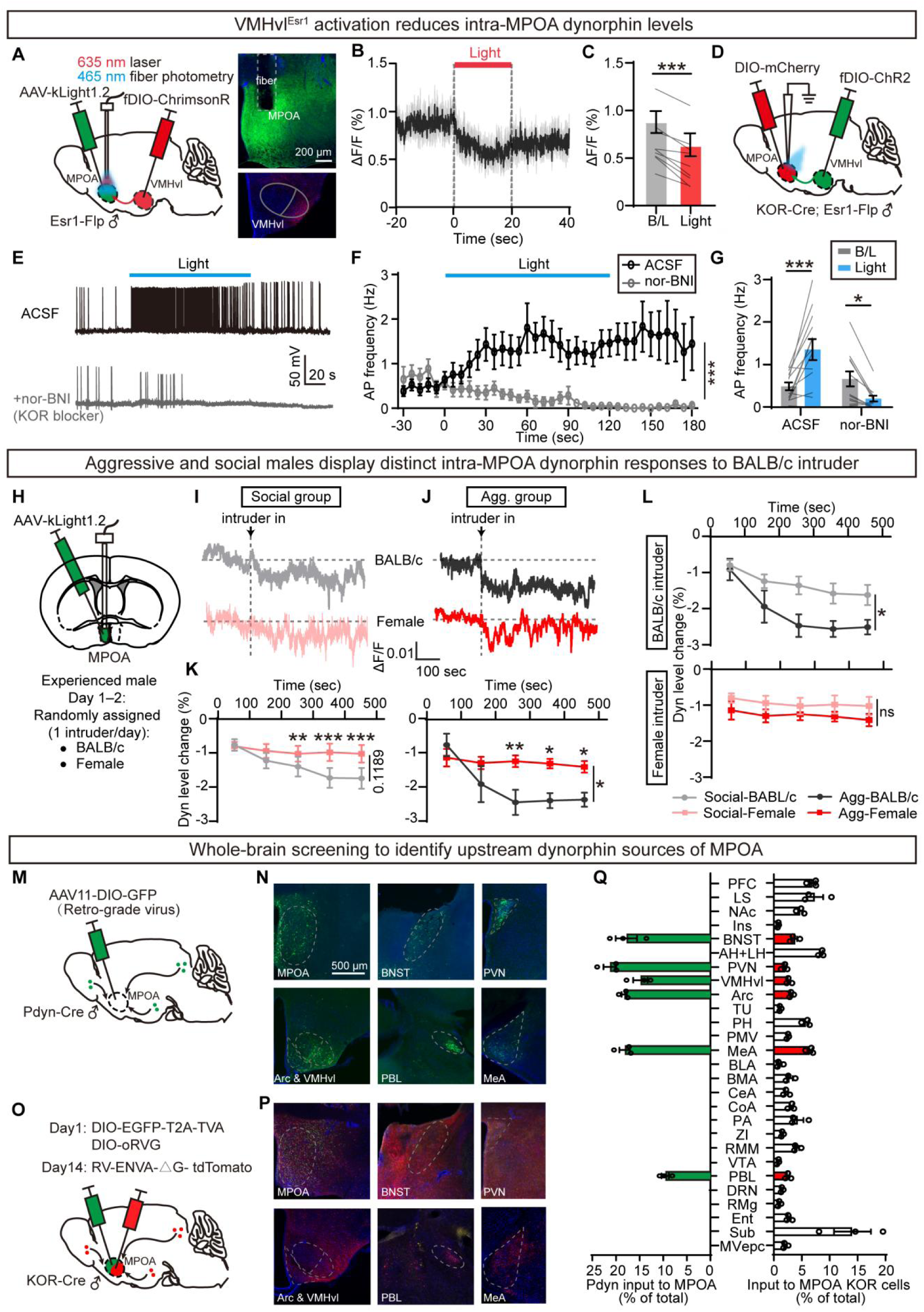
VMHvl^Esr1^ activates MPOA^KOR^ mainly via reduced dynorphin release. (A) Viral strategy and histological validation for light-activating VMHvl^Esr1^→MPOA projection and simultaneous kLight signal (dynorphin sensor) recording in the MPOA. (B) Averaged kLight signal of MPOA across animals during VMHvl^Esr1^→MPOA projection activation. (C) Mean kLight ΔF/F signal at baseline (B/L) and during light stimulation. (D) Viral strategy for synaptic recording between VMHvl^Esr1^ and MPOA^KOR^ in brain slices. (E) Representative firing traces of MPOA^KOR^ neurons during VMHvl^Esr1^→MPOA projection activation under ACSF baseline and following nor-BNI application. (F-G) The mean action potential firing frequency with 5 sec bin size (F) and frequency (G) during light in ACSF or adding nor-BNI. (H) Experimental design for kLight sensor recording during exposure to divergent social stimuli: BALB/c and female intruder stimuli. (I-J) Representative kLight signal traces from social (I) and aggressive (J) male mice exposed to BALB/c or female stimuli. (K) Quantification of kLight signal reduction across divergent social stimuli in social (left) and aggressive male (right) . (L) Quantification of kLight signal reduction in social and aggressive mice exposed to BALB/c (top) and female (bottom) stimuli. (M-N) Viral strategy (M) and histology (N) for retrograde mapping of upstream dynorphin afferent inputs to the MPOA. (O-P) Viral strategy (O) and histology (P) for monosynaptic rabies tracing of upstream inputs to MPOA^KOR^ neurons. (Q) Schematic summarizing dynorphin upstream afferent inputs to the MPOA and monosynaptic upstream inputs to MPOA^KOR^ neurons. Data are mean±SEM. (C) Two-tailed paired t test. (F) Two-way ANOVA. (G, K, L) Two-way ANOVA with Sidak’s multiple comparisons test. \**p* < 0.05, \*\**p* < 0.01, \*\*\**p* < 0.001; ns, not significant. n=animals; N=cells. (C) n=9. (F, G) N=13 MPOA^KOR^ cells for ACSF, and 12 for nor-BNI. (K, left) n=11. (K, right) n=6. (L, top) n=13 for social group; n=7 for aggression group. (L, bottom) n=11 for social group; n= 6 for aggression group. (Q) n=3. See also Figure S7.

Supporting this, optogenetic activation of all VMHvl glutamatergic neurons by injecting AAV-Vglut2-ChR2, encompassing both Esr1 and non-Esr1 populations, evoked monosynaptic EPSCs in 64% of recorded MPOA^KOR^ neurons (36/56=64%) (Fig. S7D-S7G)—a higher ratio than when targeting VMHvl^Esr1^ specifically (7/36=19%; Fig. S7F), indicating that the majority of direct glutamatergic input onto KOR neurons originates from non-Esr1 cells. Together, these findings support that VMHvl^Esr1^ neuronal activation drives MPOA^KOR^ activity predominantly through modulation of dynorphin–KOR neuropeptide signaling, with glutamatergic transmission playing a supplementary role; additional direct glutamatergic input from non-Esr1 VMHvl neurons further contributes to KOR neuron recruitment. These inputs collectively regulate MPOA^KOR^ excitability and thereby mediate the delay in mating initiation.

### Aggression and social males show distinct dynorphin responses to intruder types

We next assessed MPOA dynorphin dynamics using the kLight sensor during exposure to different intruder types (Fig. 7H). In aggressive and social males, introduction of a BALB/c male or a female both reduced dynorphin levels. The dynorphin decrease exposure to female was significantly smaller than that evoked by BALB/c in both groups (Fig. 7I-7K). This aligns with previous reports that females elicit weaker VMHvl^Esr1^ activation than males^8^, resulting in less dynorphin reduction and consequently greater disinhibition of MPOA^Esr1^ neurons under female versus BALB/c conditions. What’s more, BALB/c induced a significantly greater dynorphin decrease in aggressive males than in social males (Fig. 7L). This more pronounced BALB/c-evoked dynorphin decline in aggressive males may disinhibit MPOA^KOR^ neurons, thereby increase KOR neuronal activity and contribute to the mating delay observed in the presence of a BALB/c competitor. Together, these results demonstrate that VMHvl^Esr1^-mediated regulation of MPOA dynorphin varies both by group (aggression vs. social) and by intruder type. These differences tune MPOA^KOR^ activity and the degree of inhibition onto MPOA^Esr1^ neurons, ultimately modulating sexual behavior in male mice.

### Arc^Pdyn^ neurons are a major source of dynorphin in the MPOA

To identify upstream dynorphin sources to the MPOA, we injected the retrograde tracer AAV11-DIO-Flp-GFP into the MPOA of Pdyn-Cre males (Fig. 7M). Retrogradely labeled dynorphinergic inputs were found in multiple regions, including the bed nucleus of stria terminalis (BNST), paraventricular nucleus (PVN), VMHvl, Arc, medial amygdala (MeA), and lateral parabrachial nucleus (PBL) (Fig. 7N), which were independently confirmed by monosynaptic rabies tracing from MPOA^KOR^ neurons (Fig. 7O-7Q). To determine which region mediates the BALB/c-evoked dynorphin decline, we injected AAV11-DIO-Flp into the MPOA, followed by fDIO-GCaMP into each candidate upstream region to record activity of Pdyn neurons projecting to MPOA (Fig. 8A). Upon BALB/c exposure, Pdyn neurons in VMHvl and BNST showed robust calcium increases, whereas those in PVN, MeA, and PBL showed no response (Fig. 8B and S8A-S8B). In contrast, Arc^Pdyn^ neurons displayed a significant decrease in calcium during BALB/c contact (Fig. 8B-8C), identifying the Arc as a likely primary source of the dynorphin decline observed in the MPOA.

**Figure 8.**
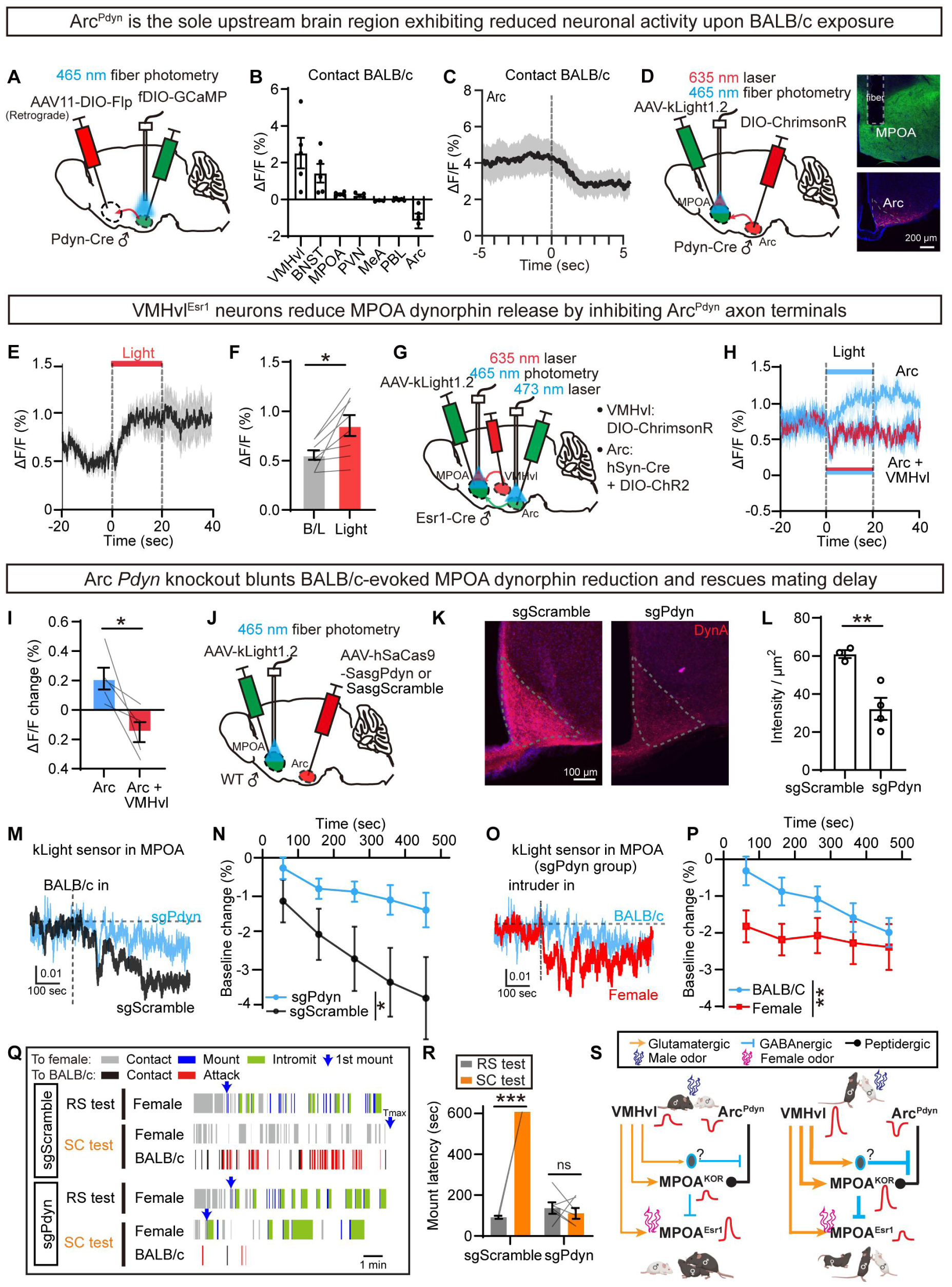
Arc^Pdyn^ neurons provide the major dynorphin source to MPOA. (A) Viral strategy for Ca^2+^ imaging of distinct dynorphin-expressing upstream inputs innervating the MPOA. (B) Mean ΔF/F of contacting BALB/c across different MPOA upstream regions. (C) Averaged PETH of ΔF/F recording in Arc^Pdyn^ neurons across animals aligned to the onset of contacting BALB/c. (D) Viral strategy and histology for light-activating Arc^Pdyn^→MPOA projection with simultaneous kLight dynorphin sensor recording in the MPOA. (E) Averaged kLight signal in the MPOA across animals during activating Arc^Pdyn^→MPOA projection. (F) The mean ΔF/F signal at baseline and during light stimulation. (G) Viral strategy and histological validation for dual optogenetic activation of Arc^Pdyn^ neurons and the VMHvl^Esr1^→MPOA projection, with simultaneous MPOA kLight recording. (H) Averaged MPOA kLight signals across animals during Arc-only activation (blue trace) and combined activation of Arc and VMHvl^Esr1^→MPOA projections (blue + red trace). (I) The mean ΔF/F signal during activating Arc only versus simultaneous activation of Arc + VMHvl^Esr1^ inputs. (J) Viral strategy for CRISPR-mediated *Pdyn* knockout in the Arc and simultaneous MPOA kLight signal recording. (K-L) Immunostaining (K) and intensity (L) of DynA immunostaining in the Arc of sgScramble and sgPdyn mice. (M) Representative MPOA kLight signal traces during BALB/c intruder exposure in Arc-targeted sgScramble and sgPdyn mice. (N) Quantification of MPOA kLight signal reduction in sgScramble and sgPdyn mice upon BALB/c intruder exposure. (O) Representative MPOA kLight signal traces in Arc sgPdyn mice during exposure to BALB/c and female intruders. (P) Quantification of kLight signal reduction in Arc sgPdyn mice following exposure to BALB/c versus female intruders. (Q) Behavioral raster plots of male mice from sgScramble and sgPdyn groups during regular sexual (RS) tests and sexual competition (SC) tests. Tmax indicates the end of the test. (R) Mount latency of the sgScramble and sgPdyn group males during RS test and SC test. (S) Schematic summary illustrating the hypothalamic circuit mechanism underlying mating delay in the sexual competition paradigm. Data are mean±SEM. (F, I) Two-tailed paired t test. (L) Two-tailed unpaired t test. (N, P) Two-way ANOVA. (R) Two-way ANOVA with Sidak’s multiple comparisons test. \**p* < 0.05, \*\**p* < 0.01, \*\*\**p* < 0.001; ns, not significant. n=animals. (B) n=5 for VMHvl, BNST, MPOA, PVN, PBL and Arc; and n=3 for MeA. (L) n=3 for sgScramble and n=4 for sgPdyn. (N) n=6 for sgScramble and n= 8 for sgPdyn. (R) n=3 for sgScramble and n=6 for sgPdyn. (F) n=8. (I) n=5. (P) n=6. See also Figure S8-S9.

To directly test whether Arc^Pdyn^ axons release dynorphin in the MPOA, we optogenetically activated the Arc^Pdyn^→MPOA projection while recording the MPOA dynorphin sensor signal (Fig. 8D). Opto-stimulation produced a significant rise in kLight signal (Fig. 8E-8F), confirming pathway-driven dynorphin release. Given that VMHvl^Esr1^→MPOA activation reduces dynorphin (Fig. 7B), we hypothesized that VMHvl^Esr1^ neurons suppress Arc^Pdyn^ terminal release. To test this, we first activated Arc alone to elevate dynorphin via ChR2, then co-activated VMHvl^Esr1^→MPOA projection using ChrimsonR (Fig. 8G). The dynorphin increase was markedly attenuated by VMHvl^Esr1^ co-activation (Fig. 8H-8I), indicating that VMHvl^Esr1^ neurons suppress Arc^Pdyn^ terminals in MPOA to reduce dynorphin release, consistent with previous finding that activation of VMHvl^Esr1^→MPOA projection induced a significant decrease in kLight signal in MPOA (Fig. 7B).

To confirm the functional contribution of Arc^Pdyn^ neurons to BALB/c-evoked dynorphin dynamics and mating delay, we knocked out *Pdyn* in the Arc using CRISPR-based AAV-Cas9-sgPdyn, with AAV-Cas9-sgScramble as control (Fig. 8J). Immunohistochemical staining for dynorphin A confirmed effective knockdown (Fig. 8K-8L). Arc *Pdyn* elimination significantly blunted the MPOA dynorphin decrease evoked by BALB/c exposure (Fig. 8M-8N), while leaving the female-evoked decrease unchanged (Fig. S9A-S9C); notably, the female-evoked decrease was then larger than the BALB/c-evoked decrease following gene knockout (Fig. 8O-8P). Finally, we examined the effect of *Pdyn* gene knockout on male mating behavior during sexual competition. Aggression group males still attacked BALB/c intruders but no longer showed a significant mating delay; BALB/c presence failed to significantly increase mount latency relative to regular sexual behavior tests (Fig. 8Q–8R). *Pdyn* knockout also significantly upregulated *Oprk1* expression in the MPOA of aggressive males (Fig. S9D–S9F), shifting it toward the higher levels characteristic of social mice (Fig. 6P–6S). Given that elevated *Oprk1* expression correlates with lower KOR neuron activity, this upregulation may inhibit mating-promoting MPOA^Esr1^ neurons, thereby relieving the mating delay. Collectively, these findings establish that Arc^Pdyn^ neurons are a key upstream source regulating BALB/c-evoked dynorphin dynamics in the MPOA and subsequent suppression of mating behavior when males face both a rival and a female partner, revealing a critical role for this VMHvl^Esr1^/Arc^Pdyn^→MPOA^KOR^→MPOA^Esr1^ circuit in governing sexual behavior under sexual competition (Fig. 8S).

## DISCUSSION

This study delineates a hierarchically organized hypothalamic dynorphin-KOR neuromodulatory circuit that governs the behavioral priority between male–male aggression and male–female reproductive behavior in male mice under sexual competition. Males with aggression training display prominent delays in mating initiation specifically when exposed to a male competitor, while social males without aggressive capability show no alterations in sexual behavior. This behavioral prioritization is mediated by two core upstream neural modules: VMHvl glutamatergic neurons and Arc dynorphinergic neuronal populations. These two distinct pathways converge onto an uncharacterized hub of KOR-expressing inhibitory interneurons in the MPOA. Glutamatergic VMHvl input provides direct excitatory drive onto MPOA^KOR^ neurons. Concurrently, given that KOR is an inhibitory Gi/o-coupled receptor, the greater reduction in Arc-derived dynorphin signaling upon exposure to a male conspecific in aggressive males relieves KOR-mediated inhibition. Together, these mechanisms recruit MPOA^KOR^ GABAergic neurons, which subsequently suppress mating-promoting MPOA^Esr1^ neurons and thereby inhibit mating behavior.

Collectively, these findings uncover a VMHvl^Esr1^/Arc^Pdyn^→MPOA^KOR^→MPOA^Esr1^ hypothalamic microcircuit that dynamically modulates reproductive behavior according to aggressive social experience, revealing a core circuit mechanism underlying sexual competition-induced reproductive behavioral plasticity.

### Behavioral Strategic Dichotomy: Resolving the “Fight or Mate” Trade-Off via Internal State and Social Context

In nature, males frequently face a behavioral trade-off: attack a rival to secure territory and mating access, or mate immediately to maximize reproductive opportunity^20^. Our findings have demonstrated that this trade-off depends on the need to establish social dominance, which varies with individuals’ aggression levels (Fig. 1B-1F), intraday aggressive experience (Fig. 1O-Q), and the type of intruders (Fig. S1F-S1I). Aggressive male mice adopt a “compete-first, mate-later” strategy—they preferentially attack competitors before mating, and this mating delay is eliminated if dominance is established prior to female introduction. Non-aggressive social males, by contrast, mount females promptly regardless of competitor presence, adopting an “avoid-conflict, mate-quickly” strategy. This phenotype-dependent allocation between aggression and mating is consistent with classic strain-comparison studies in CBA and ST mice: the more aggressive CBA males typically confront and defeat rivals before mating, whereas the less aggressive ST males, when they mate successfully, achieve this without first defeating the competitor^21^. The context-dependent reallocation between aggression and mating observed here is not limited to rodents. In Drosophila, males suppress courtship when a rival is present^22^; in chimpanzees, low-ranking males shift from direct competition to opportunistic mating^23^. Together, these findings suggest that males adopt flexible strategies for balancing aggression and mating, with their behavioral priorities depending on both social context and intrinsic aggressiveness. Evolutionarily, this behavioral divergence represents an adaptive reproductive trade-off coupled with distinct ejaculate strategies across social ranks^24^. Aggressive dominant males delay mating to establish social dominance, securing mating priority and investing in high-quality ejaculates with larger sperm reserves to produce fitter offspring. In contrast, non-aggressive subordinate males adopt fast, opportunistic mating to offset social inferiority, relying on frequent matings rather than superior sperm quality to secure reproductive success^25^.

We further ruled out trivial explanations and confirmed that this mating delay reflects an active, adaptive strategy rather than passive behavioral impairment. First, the delay is not due to reduced sexual motivation nor ability. Aggressive males exhibit robust preference for female cues and display normal consummatory mating performance under competition. Second, this delay is not induced by general aggressive arousal or sensory distraction. Males show comparable aggression toward juvenile intruders but no mating delay, confirming that only reproductively competitive adult male rivals trigger this behavioral suppression. Third, aggressive males actively suppress courtship USVs in the presence of male competitors. Importantly, competition exerts divergent effects on mating behavior and USV vocalizations: mounting behavior remains largely intact, whereas USVs are severely inhibited, suggesting that these two aspects of reproductive behavior are mediated by separable neural circuits. This USV suppression is consistent with the social audience effect observed in multi-male contexts, wherein males reduce conspicuous courtship USVs to avoid escalating conflicts with rivals^26,27^.

### MPOA Inhibitory Interneurons: The Missing Relay Bridging Aggression and Reproductive Circuits

Reciprocal interactions between the VMHvl and MPOA coordinate aggression and mating: VMHvl→MPOA signaling suppresses reproductive behavior^8^, whereas caudal MPOA→VMHvl projections restrain aggression toward a dominant rival^10^. Given that VMHvl neurons are glutamatergic, one would expect this pathway to directly excite its downstream targets. Yet how this pathway ultimately inhibits mating-promoting MPOA^Esr1^ neurons has remained unclear.

Our study resolves this longstanding puzzle by identifying MPOA^KOR^ GABAergic interneurons as the key inhibitory relay bridging VMHvl^Esr1^ and MPOA^Esr1^ circuits. Multiple orthogonal approaches establish their causal intermediary role in the aggression-mating trade-off: (1) optogenetic activation of MPOA^KOR^ interneurons is sufficient to recapitulate competitor-induced mating suppression even in the absence of rivals; (2) silencing this interneuron population completely eliminates competition-triggered mating delay without altering baseline aggression; (3) rabies tracing and ex vivo patch-clamp recordings demonstrate that these interneurons receive monosynaptic input from VMHvl neurons and send inhibitory synapses to MPOA^Esr1^ mating neurons; and (4) MPOA^KOR^ neurons are robustly disinhibited via reduced dynorphin release from VMHvl^Esr1^→MPOA terminals, an effect that is occluded by KOR antagonist pretreatment.

Consistent with high-resolution MERFISH hypothalamic profiling^13^, MPOA^KOR^ interneurons form a transcriptionally distinct inhibitory cluster that are specifically activated by male aggression, with only ∼20% overlap with mating-related MPOA^Esr1^ neurons. Our identification of the MPOA^KOR^→MPOA^Esr1^ pathway does not contradict the sparse local connectivity framework in the hypothalamus^14–16^. Rather, it complements this view by revealing a “sparse but convergent” architecture: paired recordings have shown that local chemical synapses within hypothalamic nuclei are rare^14–16^, and in the MPOA specifically, dual whole-cell recordings failed to detect inhibitory connections among neighboring neurons^16^. Yet when we synchronously activated the MPOA^KOR^ interneuron population with optogenetics, we observed robust IPSCs (∼600 pA) in MPOA^Esr1^ neurons—currents comparable to those reported for projections from MPOA to VMHvl^10^. This demonstrates that sparse inputs can pool to generate potent convergent inhibition, reconciling the low pairwise connection probability with the strong functional influence we observe. Notably, although dynorphin acts on KORs from Arc^Pdyn^ terminals rather than local sources, its regulatory role is essential for MPOA^KOR^ interneuron excitability and consequently MPOA^Esr1^ activity, reinforcing the key role of neuropeptides in hypothalamic control of social decision-making^14^. Thus, long-range peptidergic modulation and local GABAergic inhibition converge onto MPOA^Esr1^ neurons to gate reproductive behavior.

A defining feature of the dynorphin–KOR circuit is its profound experience-dependent plasticity. We found that prior social history drives divergent cell-intrinsic properties and dynorphin dynamics in MPOA^KOR^ interneurons, shaping long-term behavioral strategies. In non-aggressive males, repeated non-threatening exposure to male conspecifics induces progressive habituation of MPOA^KOR^ calcium responses, driven by gradual adaptation of dynorphin release—the initial large reduction during the first day wanes over successive encounters, progressively dampening KOR-mediated inhibition onto MPOA^Esr1^ neurons and enabling rapid mating initiation in subsequent competitive contexts. In contrast, aggressive males show persistent MPOA^KOR^ activation after repeated agonistic encounters, with sustained strong dynorphin reduction and no adaptation, maintaining potent KOR-mediated inhibition onto MPOA^Esr1^ neurons and locking in a “compete-first, mate-later” strategy.

### Glutamate and dynorphin converge in the MPOA to regulate sexual behavior in the sexual competition test

Notably, VMHvl regulates MPOA^KOR^ neuronal excitability through two pathways that endow the circuit with multi-timescale computational capacity for social decision-making (Fig. S7E–S7G and 5I). Monosynaptic retrograde tracing reveals that glutamatergic inputs to MPOA^KOR^ cells arise predominantly from non-Esr1 VMHvl^Vglut2^ neurons (Fig. S7C), which mediate fast excitation, whereas VMHvl^Esr1^ neurons contribute only ∼20% of these inputs. Despite this sparse contribution, the VMHvl^Esr1^-to-MPOA^KOR^ direct glutamatergic projection is required for the poly-synaptic IPSCs from VMHvl^Esr1^ to MPOA^Esr1^. Convergently, the VMHvl^Esr1^-to-MPOA projection serves a critical peptidergic neuromodulatory function: optogenetic activation of VMHvl^Esr1^ reduces Arc^Pdyn^-derived dynorphin levels in the MPOA (Fig. 8H), relieving constitutive KOR-mediated inhibition and persistently elevating MPOA^KOR^ excitability. Thus, through the VMHvl^Esr1^/Arc^Pdyn^→ MPOA^KOR^→MPOA^Esr1^ circuit, the rapid glutamatergic pathway and the sustained disinhibitory pathway together integrate both aggressive motivational states and immediate social signals upon encountering BALB/c mice, ultimately gating reproductive decision-making.

Arc^Pdyn^ neurons provide a prominent dynorphinergic input to MPOA^KOR^ neurons, and their activity closely tracks local peptide dynamics in the MPOA: BALB/c encounters suppress Arc^Pdyn^ calcium signals in parallel with the decline in MPOA dynorphin, whereas optogenetic activation of Arc^Pdyn^ neurons elevates dynorphin levels locally. Notably, Arc-specific *Pdyn* knockout attenuated the BALB/c-evoked reduction in MPOA dynorphin and increased MPOA *Oprk1* expression, suggesting a compensatory adjustment of the ligand–receptor system following sustained loss of Arc-derived dynorphin. This molecular shift converted the mating strategy of aggressive males toward a social-like phenotype, eliminating competition-induced mating delay without impairing aggressive motor output. By contrast, female-evoked dynorphin modulation remained intact, indicating that distinct social contexts engage separable sources or regulatory mechanisms of MPOA dynorphin—specifically, male-related signals originate from Arc, whereas female-related signals derive from a different, as-yet-unidentified brain region. Taken together, Arc^Pdyn^ signaling appears to gate how competition influences mating rather than controlling aggression itself, while coordinated plasticity of dynorphin release and KOR expression determines whether rival cues are translated into reproductive suppression.

Our results further suggest that rival-evoked dynorphin suppression is achieved through coordinated regulation of both Arc^Pdyn^ somatic activity and its terminal release within the MPOA. BALB/c exposure reduced calcium activity in Arc^Pdyn^ somata, indicating that rival cues suppress the upstream activity of dynorphin-producing neurons. Concurrently, activation of VMHvl^Esr1^ terminals in the MPOA further reduced local dynorphin signals, consistent with an additional terminal-level mechanism that gates release from Arc^Pdyn^ axons. The convergence of these two regulatory layers may produce a greater suppression of dynorphin signaling than somatic decrease alone, ensuring robust attenuation during sexual competition. However, given that VMHvl^Esr1^ neurons are glutamatergic, the precise mechanism by which their axonal projections in the MPOA reduce dynorphin release from Arc^Pdyn^ axons remains unknown.

Collectively, this dual-mode circuit design, combining glutamatergic excitation and opioid neuromodulation, enables the MPOA circuit to integrate acute social threats and sustained competitive states, supporting precise and flexible control of social decision-making.

### Arc^Pdyn^ Neurons: Repurposing a Canonical Metabolic Hub for Social Competitive Gating

The Arc nucleus is historically defined as a central hypothalamic integrator of energy homeostasis and feeding behavior^28^. Emerging work has expanded this framework, demonstrating that discrete Arc neuronal subpopulations are repurposed to modulate innate social and reproductive behaviors beyond metabolism. In males, Arc neurons are implicated in a range of social and reproductive processes, including ejaculatory reward via intra-hypothalamic circuitry^29^ and adolescent isolation-dependent social need^30^. In females, Arc^Agrp^ neurons suppress maternal nesting behavior via MPOA projections^31^ and attenuate fertility by delaying estrous cycles through direct inhibition of Kiss1 neurons^32^. Our findings further extend this paradigm by establishing Arc^Pdyn^ neurons as a key regulator specifically during male sexual competition, gating reproductive behavior through convergent dynorphin-mediated inhibition onto MPOA^Esr1^ neurons.

Intriguingly, Arc^Pdyn^ neurons share molecular and regulatory features with the well-characterized Arc^Agrp^ ensemble, supporting a unified model of the Arc as a versatile state-integration hub. Both populations converge on the MPOA as a final common behavioral control node: starvation-activated Agrp neurons suppress mating and maternal behavior, whereas competition-recruited Pdyn-KOR signaling delays male mating. Molecular profiling further links their transcriptional identities to divergent social functions: approximately 75% of Arc^Pdyn^ neurons co-express NPY, a canonical marker of Agrp neurons^33^. Recent work has extended this principle to the social domain: Agrp neurons are activated by social isolation and suppressed by reunion^30^, whereas Pdyn neurons are inhibited by direct male intruder contact—consistent with our observation that Arc^Pdyn^ activity decreases upon encountering a BALB/c intruder. Furthermore, this circuit framework may also help explain that hungry male mice mate with a female alone but prioritize feeding when both food and a female are available^34^. We speculate that under food-deprived conditions, sustained Arc^Agrp^ activity drives dynorphin release, which inhibits MPOA^KOR^ and disinhibits MPOA^Esr1^ to permit mating; food presence suppresses this pathway, allowing increased KOR tone to inhibit Esr1 and prioritize feeding. Thus, Arc^Pdyn^ neurons should not be viewed solely as endocrine regulators, but as rapid sensors of metabolic and social states that adjust reproductive strategy via KOR-mediated gating of MPOA^Esr1^. Whether metabolic state modulates Arc^Pdyn^ excitability and whether local microcircuits mediate Agrp-Pdyn crosstalk remain open questions.

#### Limitations of the study

While our work establishes the core circuit logic governing competitive reproductive prioritization, several key mechanistic and ecological questions remain unresolved, highlighting opportunities for follow-up investigation.

### First, the synaptic mechanism underlying VMHvl-mediated suppression of Arc dynorphin release remains undefined

Our data confirm that VMHvl activation reduces dynorphin efflux from Arc terminals in the MPOA, but we did not identify the anatomical pathway enabling this modulation. Plausible mechanisms include: (1) indirect inhibition of Arc^Pdyn^ neurons via VMHvl-driven activation of local MPOA inhibitory interneurons; (2) descending extra-hypothalamic pathways relaying VMHvl social threat signals to the Arc; or (3) presynaptic modulation of Arc terminal release probability via remote neuromodulatory inputs. Future anterograde tracing and cell-type-specific circuit perturbation will distinguish these models.

### Second, auxiliary dynorphinergic inputs to the MPOA likely mediate context-specific behaviors beyond sexual competition

Our tracing identified additional dynorphinergic projections from the MeA and PBL to the MPOA, which are known to regulate thermoregulation and nociception. The finding that female-triggered MPOA dynorphin reduction is unaffected by Arc *Pdyn* knockdown confirms that non-Arc dynorphin sources are selectively recruited by non-competitive social stimuli. Dissecting how distinct dynorphinergic modules are gated by sensory valence (competitive male vs. receptive female) will reveal broader principles of opioid neuromodulation in hypothalamic function.

### Third, hormonal and ecological variables that set circuit baseline excitability require systematic exploration

Our experiments were conducted under controlled laboratory conditions with age-matched, gonadally intact male mice. In natural habitats, circulating gonadal steroids, chronic stress hormones, prior social rank, fluctuating female receptivity, and predation risk dynamically tune neuronal excitability. Future work should investigate how testosterone and corticosterone modulate MPOA interneuron intrinsic properties and Arc dynorphin release, to contextualize our circuit model within ecologically relevant physiological backgrounds.

### Concluding Remarks

This study provides a comprehensive, circuit-level framework explaining how mammalian brains resolve the evolutionary trade-off between social competition and reproductive success. The VMHvl/Arc→MPOA^KOR^ interneurons→MPOA^Esr1^ mating neurons pathway acts as a neural embodiment of the “compete-first, mate-later” strategy in aggressive males, while experience-dependent MPOA interneuron plasticity enables flexible adaptive behavior in non-aggressive conspecifics. By uncovering KOR-mediated neuromodulation as the core mechanism that prioritizes aggression and delays mating, we bridge longstanding gaps between hypothalamic circuit physiology, social decision-making, and evolutionary fitness theory. The repurposing of the Arc, an ancient metabolic hub, for competitive social gating further underscores the remarkable modularity and functional plasticity of hypothalamic neural circuits, which have evolved to integrate internal physiological states and external social cues to optimize behavioral output across dynamic natural environments.

## RESOURCE AVAILABILITY

### Lead contact

Further information and requests for resources and reagents should be directed to and will be fulfilled by the lead contact, Dr. Luping Yin.

### Materials availability

The newly generated AAV constructs described in this study are available from the lead contact upon request and with a completed Materials Transfer Agreement.

### Data and code availability

- All data reported in this paper will be shared by the lead contact upon request.
- This paper does not report original code. All MATLAB codes used in this manuscript are available from the lead contact upon request.
- Any additional information required to reanalyze the data reported in this paper is available from the lead contact upon request.

## ACKNOWLEDGMENTS

We thank all members of the Yin laboratory for discussions, Y.Y. Liu for genotyping assistance, and the following investigators for sharing transgenic mouse lines: B. Zhang (Westlake University) for Fos-CreER; W.S. Wang (Fudan University) for CCK-Cre; M. He (Fudan University) for H2B-GFP; X.H. Xu (Fudan University) for Esr1-Cre; Y.G. Sun for Tacr1-Flp; and Q.F. Ma (Westlake University) for KOR-Cre and Pdyn-Cre.

## AUTHOR CONTRIBUTIONS

L.Y. conceived the project, supervised the project, designed experiments, and wrote the manuscript. Q.L. co-designed experiments, conducted nearly all the experiments, analyzed the data, and co-wrote the manuscript. Y.Z. assisted with behavioral experiments and histology. X.M. assisted with Matlab code for USV analysis.

## DECLARATION OF GENERATIVE AI AND AI-ASSISTED TECHNOLOGIES IN THE WRITING PROCESS

During the preparation of this work, the authors used Doubao, DeepSeek and ChatGPT for grammar and spelling checks. After using this tool, the authors reviewed and edited the content as needed and take full responsibility for the content of the published article. No generative AI tools were used to produce any original ideas or content.

## SUPPLEMENTAL INFORMATION

Document S1. Supplemental Figures S1–S9

Data S1. Detailed statistics of all figure panels

## STAR METHODS

### EXPERIMENTAL MODEL AND STUDY PARTICIPANT DETAILS

#### Animals

All experimental procedures were approved by the Institutional Animal Care and Use Committee of the School of Life Sciences at Westlake University. This study used 7- to 12-week-old male mice of the following strains: C57BL/6J, BALB/c, Vglut2-Flp, Vgat-Flp, CCK-Cre, H2B-GFP, Fos-CreER, Esr1-Cre, KOR-Cre, Pdyn-Cre and Esr1-Flp. The Tacr1-Flp line was generated by Dr. Y. G. Sun at the Institute of Neuroscience, Shanghai^35^. H2B-GFP was originally generated from Dr. Z. Josh Huang lab from Duke university^36^. Vglut2-Flp (030212), Vgat-Flp (029591), CCk-Cre (012706), Fos-CreER (021882), Esr1-Cre (017911), KOR-Cre (035045), Pdyn-Cre (027958), and Esr1-Flp (036028) were originally obtained from the Jackson Laboratory. C57BL/6J mice were sourced from the Laboratory Animal Resources Center of Westlake University, and BALB/c mice were obtained from Charles River, China. All mouse strains were bred and maintained at the Animal Experimental Center of Westlake University. To generate double transgenic mice, we crossed KOR-Cre with Esr1-Flp mice to produce KOR-Cre;Esr1-Flp offspring, Esr1-Cre with Tacr1-Flp mice to produce Esr1-Cre;Tacr1-Flp offspring, and CCK-Cre with H2B-GFP to produce CCK-Cre;H2B-GFP offspring to label CCK+ neurons. All animals were housed in a temperature-controlled environment (22–24 °C, 55–65% humidity) under a 12-hour light/dark cycle (lights on at 07:00 a.m.) with ad libitum access to food and water.

#### Viruses

All viral vectors were purchased from commercial sources or custom-packaged as follows. AAV11-hSyn-fDIO-Cre-EGFP, AAV11-EF1a-DIO-Flp-EGFP, and AAV11-EF1a-DIO-Flp were purchased from BrainCase (Shenzhen, China). AAV2/5-hSyn-DIO-ChrimsonR-mCherry, AAV2/5-EF1α-DIO-hChR2(H134R)-EYFP, AAV2/9-hSyn-DIO-mCherry, AAV2/9-hSyn-fDIO-mCherry, AAV2/9-EF1a-fDIO-hChR2(H134R)-EYFP, AAV2/9-EF1a-fDIO-ChrimsonR-mCherry, AAV2/9-hSyn-DIO-GtACR2-P2A-EGFP, AAV2/9-EF1a-kLight1.2, AAV2/9-VGLUT2-hChR2(H134R)-EGFP, AAV2/5-EF1α-DIO-H2B-EGFP-T2A-TVA, AAV2/5-EF1α-DIO-oRVG, AAV-CMV-NLS-hSaCas9-NLS-3XHA-bGH-polyA-U6-SasgRNA1(scramble)-U6-Sasg RNA2(scramble, GTGTAGTTCGACCATTCGTG), and RV-CVS-ENVA-N2C(△G)-tdTomato were purchased from BrainVTA (Wuhan, China). AAV2/9-CAG-FLEX-GCaMP6s and AAV2/9-hEF1a-DIO-hChR2(H134R)-mCherry were purchased from Taitool (Shanghai, China). The AAV2/9-Ef1a-fDIO-GCaMP6s plasmid was first obtained from Addgene (cat. #105714) and then custom-packaged at Taitool (Shanghai, China). AAV-CMV-NLS-hSaCas9-NLS-3XHA-bGH-polyA-U6-SasgRNA1(Pdyn, CCAGCAGCTTTGGCAACGGAA)-U6-SasgRNA2(Pdyn, GCCACGGAGCCCAGAGACCGT) was custom-packaged at BrainVTA (Wuhan, China). All AAV was used at a titer of ∼5 × 10¹² vg/mL. Rabies virus was used at a titer of 2.00 × 10⁸ IFU/mL.

### METHOD DETAILS

#### Stereotaxic surgery

Mice were anesthetized with isoflurane (∼1.2%) in oxygen-enriched air and placed on a stereotaxic frame. All stereotaxic instruments and accessories (including glass capillaries and a pressure injector) were purchased from RWD (Shenzhen, China). Typical injection volumes were 30–200 nL, depending on the target area volume and virus serotype. The flow rate was adjusted accordingly to achieve a total injection duration of approximately 10 minutes, followed by a 10 minutes incubation period. Coordinates (anterior-posterior, AP; medial-lateral, ML; dorsal-ventral, DV) for injection sites were referenced from “The Mouse Brain in Stereotaxic Coordinates” by Paxinos and Franklin: MPOA (AP: 0 mm, ML: ±0.3 mm, DV: −4.95 mm), VMHvl (AP: −1.8 mm, ML: ±0.75 mm, DV: −5.6 mm), Arc (AP: −1.50 mm, ML: ±0.20 mm, DV: −5.90 mm), BNST (AP: −0.45 mm, ML: ±0.9 mm, DV: −3.6 mm), PVN (AP: −0.6 mm, ML: ±0.3 mm, DV: −4.3 mm), MeA (AP: −2.0 mm, ML: ±2.25 mm, DV: −4.6 mm), and PBL (AP: −5.4 mm, ML: ±1.2 mm, DV: −3.35 mm).

For viruses that were injected mixedly, a 1:1 ratio was used unless otherwise stated. Mice were tested three weeks after virus injection. For specialized tracing, such as rabies tracing, a mixture of AAV2/5-EF1α-DIO-H2B-EGFP-T2A-TVA and AAV2/5-EF1α-DIO-oRVG (1:1 ratio) was first injected into the MPOA. Two weeks later, the rabies virus (RV-CVS-ENVA-N2C(ΔG)-tdTomato) was injected into the same region. Mice were sacrificed ten days after the rabies virus injection.

An optic fiber (400 μm for MPOA and 200 μm for other brain regions) was implanted unilaterally above the target region (Inper, Hangzhou, China). The fiber tip was positioned 400 μm above the target for optogenetics (NA=0.37) and 150 μm for photometry (NA=0.5). The fiber was fixed with adhesive and dental cement, and mice were returned to their home cages after the cement dried.

For cannula implantation, a bilateral guide cannula (OD=0.48, CC=0.6, B=7.8, M=3.5 in mm; cat. #62022), double dummy cannula (OD=0.3, CC=0.6, G2=0 in mm; cat. #62122), and double injector (OD=0.30, CC=0.7, G1=0 in mm; cat. #62250) were purchased from RWD (Shenzhen, China). The guide cannula and dummy were implanted 0.3–0.4 mm above the MPOA. Tissue adhesive and dental cement were applied to the skull surface to fix the cannula.

#### Viral histology and immunohistochemistry

To visualize viral expression and fiber endings, the headpost was not removed until embedding in OCT. Only mice with correct viral expression and fiber tract placement were used for quantification of behavioral comparisons. For male–male interaction-induced Fos staining, mice were single-housed for over 3 days, after which a BALB/c male mouse was introduced for 15 minutes. The experimental male mice were then sacrificed 1 hour after removal of the BALB/c intruder. For the immunohistochemical staining for dynorphin A and verification of *Pdyn* knockout efficiency, mice received an intraperitoneal (i.p.) injection of colchicine (75 μg/kg) 48 hours prior to perfusion, to block axonal transport and allow dynorphin to accumulate within neuronal soma^37^. Mice were anesthetized with an overdose of isoflurane and perfused transcardially with 1× PBS at room temperature (RT), followed by ice-cold 4% paraformaldehyde (PFA) in 1× PBS. Brains were extracted and post-fixed in 4% PFA overnight at 4 °C, followed by immersion in 30% sucrose/PBS for 24–72 hours at 4 °C. Brains were sectioned at a thickness of 50 μm for immunohistochemistry.

For immunohistochemistry, brain sections were rinsed with PBS three times for 5 minutes each, and then permeabilized in 0.3% PBST (0.3% Triton X-100 in PBS) for 1 hour at RT. Sections were blocked in 10% BSA (10% bovine serum albumin in 0.3% PBST, Sigma-Aldrich, cat. #V900933) for 1 hour at RT. Slices were subsequently incubated in primary antibody solution at 4 °C overnight. The primary antibodies used were as follows: anti-Fos (1:5000, SYSY, rabbit, cat. # 226008 or guinea pig, cat. # 226308), anti-Esr1 (1:10000, rabbit, Millipore, cat. #06-935), anti-Dynorphin A (1:1000, rabbit, BMA-Biomedicals, cat. #T-4268), anti-mCherry (1:1000, rabbit, Abcam, cat. #ab183628), anti-TH (1:1000, chicken, Abcam, cat. #ab76442) and anti-HA (1:1000, mouse, Sino Biological, cat. #100028-MM10). Sections were rinsed in PBST three times and incubated in 10% BSA containing secondary antibody (1:1000, Invitrogen) for 2 hour at RT. For Esr1 immunohistochemistry, to enhance the signal, after primary antibody incubation and rinsing three times, sections were incubated in Goat anti-Rabbit Poly HRP (Thermo Fisher, cat. #B40962) for 1 hour at RT, followed by incubation in TSA Vivid Fluorophore (1:1000 in Multiplex TSA Buffer, ACD, cat. #PG-323273) for 40 minutes at 40 °C. After rinsing in PBST three times, the sections were mounted on gelatin-coated glass slides, covered with anti-fade mounting medium (Abcam, cat. #ab104135) containing DAPI (Roche, cat. #10236276001) and sealed with coverslips. Slides were scanned using an epifluorescence microscope (3D HISTECH Pannoramic MIDI II, Hungary) with a 10× objective lens.

#### RNAscope

To examine the co-labeling of TRAP (mCherry) and *Oprk1* in Fig. 3, mice were single-housed for over 3 days, after which a BALB/c male mouse was introduced into the home cage for 15 minutes to allow for attack and contact. To examine the co-labeling of *Fos* (induced by ChR2 activation) and *Oprk1* in the experiment shown in Fig. 5, mice were single-housed for over 3 days, and 475 nm blue light (1–4 mW, 20 Hz, 20 ms/pulse) was delivered unilaterally into the MPOA for 10 minutes. To examine *Oprk1* expression levels in social interaction or aggression groups in Fig. 6, mice were allowed to contact or attack an intruder BALB/c male mouse in their home cage for 15 minutes per day, lasting for 5 consecutive days.

Brains were then freshly dissected 30 minutes after the behavioral or light stimulation, frozen immediately in OCT, and sectioned at 16 μm thickness on a cryostat. Slides were stored at –80°C for up to 2 weeks and dried at −20°C for 1 hour immediately before the experiment. Slides were processed according to the RNAscope Multiplex Fluorescent v2 Assay protocol (ACD) using the Fluorescent Multiplex Reagent Kit (cat. #323100) and probes for *mCherry* (cat. #431201-C3), *Oprk1* (cat. #316111-C1, encodes the kappa opioid receptor), *Slc17a6* (cat. #1247881-C2), *Slc32a1* (cat. #319191-C3), and *Fos* (cat. #316921-C1/C3). Briefly,

sections were fixed in 4% PFA (in 1× PBS) for 10 minutes at 4 °C, then dehydrated in 50%, 70%, and 100% ethanol for 5 minutes each at RT. Next, Protease IV was applied to the slides and incubated for 10 minutes at RT. After washing with PBS, the mixed probes were applied to entirely cover each slide and incubated for 2 hours at 40 °C. The slides were then sequentially incubated with AMP1, AMP2, and AMP3 for 30 minutes each at 40 °C (except AMP3, which was 15 minutes). Using the C1 probe as an example, the slides were incubated with HRP-C1 for 15 minutes at 40 °C, followed by diluted fluorophore (1:1500 in Multiplex TSA Buffer) to label the C1 probe. Before adding the next probe, enough HRP blocker was applied to entirely cover each slide. Finally, the sections were mounted and covered with anti-fade mounting medium (Abcam) containing DAPI (Roche) and sealed with coverslips.

RNAscope data analysis was quantified using 3D HISTECH proprietary software. The number of probe copies in clustered signals was calculated manually. *Oprk1* signal counts were quantified for each cell and used for classification into the following categories: bin 0 (0 puncta), bin 1 (1–3 puncta/cell), bin 2 (4–9 puncta/cell), bin 3 (10–15 puncta/cell), or bin 4 (>15 puncta/cell). The number of DAPI-positive nuclei was calculated automatically using ImageJ, given the overwhelmingly high number of DAPI signals. A histogram was used to represent the expression level distribution, calculated as an H-score. The overall H-score was calculated by totaling the percentage of cells in each bin according to the weighted formula: H-score = 0 × (% of cells in bin 0) + 1 × (% of cells in bin 1) + 2 × (% of cells in bin 2) + 3 × (% of cells in bin 3) + 4 × (% of cells in bin 4).

#### TRAP induction

To specifically label and manipulate neurons activated during male–male attack or social contact, Fos-CreER mice received bilateral MPOA injections of AAV-DIO-ChR2-mCherry. For activity-dependent neuronal tagging, 4-hydroxytamoxifen (4-OHT; Sigma-Aldrich, cat. #H6278) was freshly prepared by dissolving 10 mg 4-OHT in 200 μL DMSO at RT, followed by dilution with 800 μl oil vehicle (castor oil:sunflower oil = 1:4, v/v) to a final concentration of 10 mg/mL. The working 4-OHT solution was administered via i.p. injection at a dose of 50 mg/kg. Three weeks after viral injection, mice were socially isolated for 3 days prior to behavioral tagging. Mice then underwent a 5-day behavioral paradigm (15 minutes daily session of male–male attack or social contact with BALB/c male conspecifics), and 4-OHT was injected immediately after the final session to permanently label neurons activated during this behavioral procedure. Two weeks after 4-OHT administration, optogenetic manipulation of Fos-tagged neurons was conducted with interleaved light and sham trials. For light trials, 473 nm blue light was delivered at 1–4 mW power, 20 ms pulse duration, and 20 Hz frequency for 120 s or 20 s per trial; identical laser parameters were used for sham trials. Light/sham stimulation was triggered upon the mice’s initiation of mounting attempts toward receptive females or social contact with BALB/c male intruders.

#### Electrophysiology

Mice were singly housed before sacrifice unless otherwise noted. For Fig. 6, mice were allowed to contact or attack intruder BALB/c in homecage for 15 minutes/day over 5 consecutive days, then sacrificed the next day for slice recording. Brains were rapidly removed in ice-cold cutting solution containing (in mM): 2.5 KCl, 1.25 NaH₂PO₄, 25 NaHCO₃, 25 glucose, 7 MgCl₂, 0.5 CaCl₂, 110 choline chloride, 11.6 sodium ascorbate, 3.1 sodium pyruvate. Slices (275 μm) were cut on Vibratome (Leica VT1200S), recovered at 36 °C for 30 minutes in ACSF (in mM: 125 NaCl, 2.5 KCl, 1.25 NaH₂PO₄, 25 NaHCO₃, 11 glucose, 1 MgCl₂, 2 CaCl₂), then kept at RT until recording. All solutions were oxygenated with 95% O₂/5% CO₂. During recording, slices were perfused with ACSF (1.5 ml/min) at 35 ℃, and MPOA fluorescent neurons were visualized under 40× for recording. Recordings used Multiclamp 700B, Digidata 1550B, and pClamp11; access resistance was monitored and cells with >25 MΩ or ≥20% change were excluded.

For voltage-clamp EPSC/IPSC recording, pipettes (4–6 MΩ) were filled with internal cesium solution (in mM: 135 CsMeSO₃, 1 EGTA, 10 HEPES, 3.3 QX-314, 4 MgATP, 0.3 Na₃GTP, 8 Na phosphocreatine, pH 7.2–7.3, 305 mOsm). Light-evoked responses were induced by 473 nm, 20 ms laser; EPSCs recorded at −70 mV and IPSCs at 0 mV. To test direct synaptic connections, 1 μM TTX (Chengdu Must Bio-technology, cat. #A0224) and 100 μM 4-AP (Sigma-Aldrich, cat. #A78403) were added; GABAergic/glycinergic transmission tested with 10 μM Gabazine (R&D, cat. #SR95531) or 5 μM strychnine (GlpBio, cat. #GD21200); The effect of dynorphin was examined using 1 μM DynA (MCE, cat. #HY-P1333). sEPSC/sIPSC frequency and amplitude were quantified using custom MATLAB code (peak–trough difference) after traces were filtered at 2 kHz and baseline-adjusted in Clampfit. For current-clamp action potential recording, pipettes were filled with internal potassium solution (in mM: 140 K-gluconate, 0.2 EGTA, 10 HEPES, 2 MgATP, 2 Na₂ATP, 3 KCl, pH 7.2–7.3, 305 mOsm). F–I curve was generated by 500-ms depolarizing steps from −20 to 360 pA (20 pA increments). To test VMHvl^Esr1^ activation of MPOA^KOR^ neurons, blue light was applied for 2 minutes; 0.2 μM nor-BNI (GlpBio, cat. #GC13645) was added to block the effects of dynorphin.

#### Behavioral tests (with USV recording)

All behavioral experiments were performed in mouse home cages under red lighting. Both overhead and side views were filmed at 25 Hz using a TTL-triggered camera. Ultrasonic vocalizations (USVs) were recorded at a 300-kHz sampling rate using an Avisoft-UltraSoundGate 116H kit with a condenser ultrasound microphone (CM16/CMPA, Avisoft-Bioacoustics) positioned 10 cm above the home cage. Audio recording was synchronized with video via a bifurcated TTL pulse generated by the TDT system and sent simultaneously to both the camera and the Avisoft interface.

#### Hormone Priming

Intruder female mice were typically group-housed. Adult females underwent ovariectomy (OVX) and were subsequently primed to estrus via hormone administration. After OVX, females recovered for at least one week. Estrus was induced as described previously^38,39^, with subcutaneous injections of 150 μL of 0.1 mg/mL estradiol (Sigma-Aldrich, cat. #E8875, dissolved in corn oil) on day −2, 75 μL estradiol on day −1, and 150 μL of 1 mg/mL progesterone (Sigma-Aldrich, cat. #P0130, dissolved in corn oil) on day 0 (the day of testing). Four hours after the final injection, receptive females were screened by introducing them into the home cage of an individually housed, sexually experienced male for 3–5 minutes; only receptive females were used as intruders.

#### Sexual competition test

Sexually experienced male mice were housed individually for at least 3 days and habituated to the recording setup in their home cage for 5 minutes before testing. On day 0, a hormonally primed receptive female was introduced into the male’s home cage for 10 minutes to conduct the regular sexual behavior test (RS test). On days 1–5, a BALB/c male (a weaker strain) was introduced into the same cage daily for 15 minutes (training session). Resident mice that attacked the intruder were assigned to the aggression group; those that did not attack were assigned to the social group. On day 6, a receptive female and a BALB/c male were simultaneously introduced into the resident’s home cage for 10 minutes (sexual competition test, SC test). A similar paradigm was used in Fig. S1E, except that the BALB/c male was replaced with a juvenile male (P15–P20).

After behavior tests, we manually annotated the following behaviors: contact, attempt to mount, mount, intromit, ejaculate, and attack. ’Contact’ was defined as active nose and paws contacting any part of the intruder’s body. ’Attempt to mount’ was characterized by the male extending its forelimbs to grasp the female’s flanks. ’Mount’ referred to the period when the male was on top of the female and held the female’s lower back with its forelimbs. ’Intromit’ was defined as deep rhythmic thrusting following mounting. ’Ejaculate’ was identified when the male stopped deep thrusting for a few seconds while continuously clutching the female, then slumped to the side. ’Attack’ was defined as a suite of actions including pushes, lunges, bites, tumbling, and fast locomotion episodes between such movements. After annotation, we analyzed the latency and duration of each behavior. Mount latency was defined as the interval between intruder introduction and the first mount. Ejaculatory latency was defined as the interval between intruder introduction and ejaculation. Mount or ejaculatory latency was recorded as 600 s if the corresponding behavior did not occur during the 10-min test. Intravaginal ejaculation latency time (IELT) was defined as the interval between the first intromission and ejaculation. We also recorded USVs produced by the male to attract the female. USV syllables were manually counted by custom-written MATLAB code based on the following criteria: frequency ranging from 30 to 125 kHz, defined as a continuous occurrence longer than 8 ms per bout, with an inter-bout interval of at least 16 ms.

#### Three chamber social preference test

The test arena (L×W×H: 50 cm ×18 cm×38 cm) was evenly divided into three compartments with a 6-cm wide opening between the neighboring compartments. All test and stimulus animals were habituated to the chamber for at least two days, 30 minutes each day at non-overlapping time. To test male’s social preference between female and BALB/c male, on the day of testing, a hormonal primed female (4-6 hours after progesterone injection) and a BALB/c male was each placed under a wire cup positioned in one of the side compartments. Then, the testing male was introduced into the middle compartment, and allowed to explore freely for 10 minutes. A female-preference index was calculated by dividing the differential time the test male spent on investigating the female and BALB/c chamber by the total time spent on investigating both chambers. A positive value indicates preference directed toward the stimulus female, whereas a negative value indicates preference directed toward BALB/c.

#### Behavioral optogenetics

Three weeks after virus expression, mice with implanted optic fibers were habituated to the testing environment. Fibers were connected to a TDT laser generator via a 0.37 NA patch cord. For ChR2-mediated optogenetic activation, 475 nm light (1–4 mW, 20 ms pulses, 20 Hz) was delivered for 2 minutes upon the male’s first mount attempt; sham stimulation served as control. For combined optogenetics and fiber photometry, 620 nm red light (1–4 mW, 20 ms pulses, 20 Hz) was used to activate ChrimsonR, while 465 nm blue light was used for GCaMP photometry. For GtACR2-mediated opto-inhibition, constant 475 nm blue light was applied for 10 minutes, immediately after which a female and a BALB/c male were simultaneously introduced for the sexual competition test.

#### Cannula drug local infusion

Sexually experienced male mice with cannula implantation were tested 1 week after surgery for full recovery. Mice underwent the sexual competition test and regular sexual test on separate days. To activate KOR receptors, 250 nL per side of 0.08 µM dynorphin A was injected into the MPOA through the cannula using an injector (RWD, cat. #62250) 10 minutes before testing, during which the animal was head-fixed on a running wheel, as previously reported^19^; control animals received saline and underwent the same tests. During the post-injection waiting period, animals were returned to their home cages. To block KOR receptors, the antagonist nor-BNI (7.5 mM, 250 nL per side) was administered via cannula; saline controls were tested on the day prior to nor-BNI testing. Given that nor-BNI is a long-acting inhibitor, based on previous reports^40^, behavioral tests were performed 24 hours after injection.

#### Fiber photometry

Three weeks after surgery and optic fiber implantation, calcium signals were recorded using a fiber photometry system (TDT). A fiber patch cord (Inper) connected to the system was attached to the implanted optic fiber. A 465 nm LED light source (30 μW) excited GCaMP fluorescence through the patch cord and implanted fiber. The instantaneous ΔF/F was calculated as (F_raw_ – F_baseline_) / F_baseline_. The peak ΔF/F was defined as the maximum ΔF/F during a behavior minus the baseline (mean ΔF/F from –5 to –1 s preceding the behavior). For dynorphin sensor recordings, the baseline was defined as the average filtered F from –300 to 0 s before intruder introduction. The change rate during a period was calculated as (average filtered F in that period – baseline) / baseline; for example, change rate (0–100 s) = [F (0–100 s) – F (–300 to 0 s)] / F (–300 to 0 s).

For combined photometry and optogenetics, mice received light stimulation in their single-housed home cage. The mean ΔF/F during a behavior was calculated as the average during light or sham stimulation minus the mean normalized ΔF/F from –20 to 0 s preceding the light/sham. To block GABA_A_ receptors during VMHvl^Esr1^ activation in Fig. 2F, 0.75 mg/kg bicuculline (MCE, cat. #HY-N0219) was administered i.p. 1 hour before recording. To activate KOR during VMHvl^Esr1^ stimulation in Fig. 5C, 5 mg/kg U50488 (Sigma-Aldrich, cat. #U111) was administered i.p. 1 hour before recording.

#### Statistics

All plots and graphs were edited by Adobe Illustrator for final publication. All statistical analyses were performed using MATLAB or Prism 9 and were two-tailed. Parametric tests, including paired t-test, unpaired t-test, and one-way ANOVA, were used when distributions passed the Shapiro–Wilk normality test (except for two-way ANOVA, where data distribution was assumed to be normal); otherwise, nonparametric tests were applied. Statistical significance between two paired groups was determined using paired t-test (normal data), or Wilcoxon matched-pairs signed rank test (non-normal data). For two unpaired groups, unpaired t-test (normal data, equal variance), Welch’s unpaired t-test (normal data, unequal variance), or Mann-Whitney test (non-normal data) was used. For repeated measures across more than two groups, RM one-way ANOVA with Tukey’s multiple comparisons test (normal data) or Friedman test with Dunn’s multiple comparisons test (non-normal data) was used. Differences between two groups were analyzed using two-way ANOVA with Sidak’s multiple comparisons test, and differences among more than two groups were analyzed using two-way ANOVA with Tukey’s multiple comparisons test. Throughout this study, n denotes the number of animals and N denotes the number of cells. \**p* < 0.05; \*\**p* < 0.01; \*\*\**p* < 0.001.

